# Decoding the Protein Composition of Whole Nucleosomes with Nuc-MS

**DOI:** 10.1101/2020.09.08.287656

**Authors:** Luis F. Schachner, Kevin Jooβ, Marc A. Morgan, Andrea Piunti, Matthew J. Meiners, Alexander Lee, Jared O. Kafader, Marta Iwanaszko, Marcus A. Cheek, Jonathan M. Burg, Sarah A. Howard, Michael-Christopher Keogh, Ali Shilatifard, Neil L. Kelleher

## Abstract

Nuc-MS characterizes histone modifications and variants directly from intact endogenous nucleosomes. Preserving whole nucleosome particles enables precise interrogation of their protein content, as for H3.3-containing nucleosomes which had 6-fold co-enrichment of variant H2A.Z over bulk chromatin. Nuc-MS, validated by ChIP-seq, showed co-occurrence of oncogenic H3.3K27M with euchromatic marks (e.g., H4K16ac and >15-fold enrichment of H3K79me2). By capturing the entire epigenetic landscape, Nuc-MS provides a new, quantitative readout of nucleosome-level biology.

---

The post-translational modifications (PTMs) decorating the four core histones in an intact nucleosome encode information for nuclear effectors that trigger defined cellular events critical to health and disease.^1–4^ For decades, affinity reagents, mass spectrometry and proteomics have played key roles in characterizing the many histone isoforms and PTMs involved in epigenetic regulation.^5–7^ However, the standard practice of relying on digestion^8^ and/or denaturation^5^ removes the linkage between modifications and their nucleosomes of origin (**Fig. 1a, at left**). By digesting histone mixtures into small peptides, correlations among PTMs are forfeited, precluding strong assertions about the original composition of the intact histone. Similarly, when a nucleosome population is denatured, information about co-localization of histone isoforms and PTMs within the same nucleosome particle is lost. This inference problem diminishes our understanding of the organization and impact of co-occurring modifications, isoforms and mutations in epigenetics and disease pathogenesis.^9, 10^

**Figure 1.**
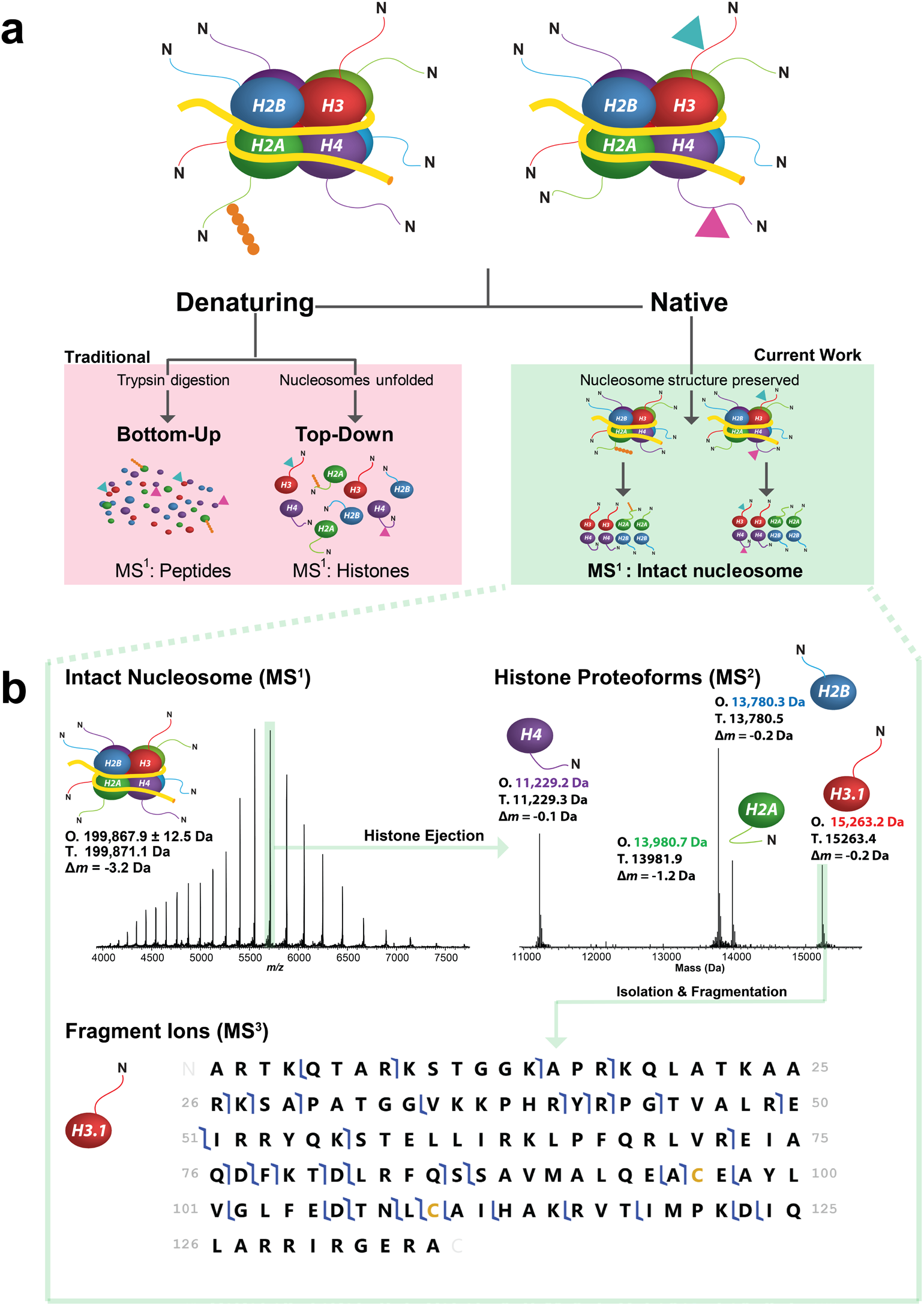
Three different strategies of histone analysis, including the new approach for direct interrogation of intact nucleosomes. **(a)** Three MS workflows for analyzing a hypothetical nucleosome mix: bottom-up, denaturing top-down, and Nuc-MS (based on native electrospray). Current methods use either protease-derived peptides or whole histones under denaturing conditions to detect histone PTMs **(a, left side)** and are thus blind to the modification state of the intact nucleosomes; these approaches cannot differentiate the two hypothetical nucleosomes. In contrast, Nuc-MS employs native electrospray to read out PTMs directly from the intact nucleosome **(a, right side)**. **(b) Nuc-MS performed** in three steps on an intact unmodified nucleosome. **First**, the mass of intact nucleosomes is measured (MS^1^: O, observed average mass; T, theoretical mass; Δ*m*, error). **Second**, a single nucleosome charge state (e.g. 35+ ions highlighted in green) is isolated and activated by collisions with nitrogen to eject all intact histones (MS^2^, reporting monoisotopic masses). **Third**, each histone is isolated and further activated to create backbone fragmentation products that characterize the proteoforms (MS^3^; blue flags indicate fragment ions matching uniquely to human histone H3.1, depicted as a graphical fragment map at bottom. Green highlight in the upper right of panel reflects the intact precursor).

To close these knowledge gaps and replace inference with direct measurement, we developed Nuc-MS, a method based on native electrospray that neither digests nor denatures nucleosomes^5^ and consists of three stages of tandem MS^11^ that capture the entire protein composition of intact nucleosomes.

We begin with synthetic nucleosomes, which have been previously subjected to intact mass analysis to assess the mechanism of assembly^12^ and structural stability.^13^ Here, we use these highly defined samples to establish proof-of-concept and benchmark Nuc-MS, which combines native mass spectrometry with direct nucleosome fragmentation by tandem MS (**Fig. 1a, far right**).

For unmodified nucleosomes containing 108.6 kDa of protein and 91.3 kDa of DNA, the Nuc-MS process is shown in **Figure 1b**. In brief, intact nucleosomes were measured with a 3.1 Da error. Isolation and activation of the 35+ charge state of the nucleosome using high-energy collisional fragmentation (HCD) resulted in the ejection of core histones, whose intact masses were detected with isotopic resolution in the same spectrum (**Fig. 1b, MS**^2^). Additional details of the three-stage process are provided in **Supplemental Figs. 1, 2,** and **3**. With this first report of controlled disassembly of nucleosome core particles, we next asserted the quantitative nature of Nuc-MS by mixing two synthetic nucleosomes bearing H3K36me1 and H3K36me2 in a 1:1 ratio. Readout of this sample using Nuc-MS showed a ratio of 49.2 ±2.5% and 50.8 ±3.3% (n = 3) for the integrated areas of ejected H3K36me1 and H3K36me2 proteoforms, respectively (**Supplemental Fig. 4**).

**Figure 2.**
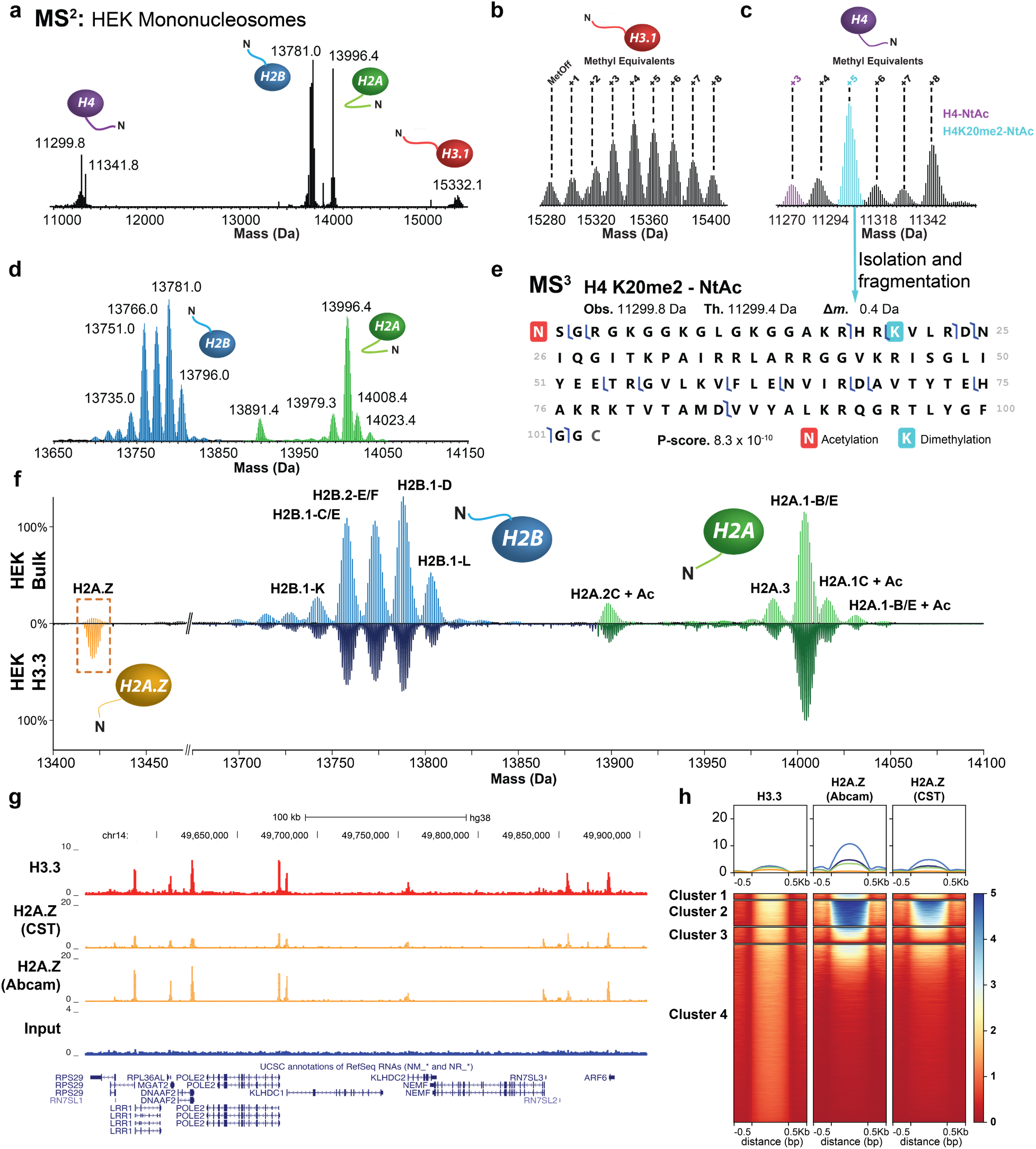
Nuc-MS analysis of endogenous mononucleosomes from HEK. **(a)** MS^2^ spectrum of ejected histones (reported as deconvoluted, average masses), demonstrating detection of all core histones and their proteoform distributions from fragmented nucleosomes in the range of 6000-9000 *m/z* (average of n = 3). **(b-d)** Spectral regions in the mass domain containing the isotopic distributions for the proteoforms of histones H3.1, H4, H2B and H2A proteoforms. **(e)** The proteoform highlighted in cyan in (e) was isolated and fragmented. The fragmentation data were then used to produce the fragmentation map shown, thus characterizing the proteoform as N-terminally acetylated H4K20me2. **(f)** Comparison of H2A and H2B proteoform profiles in HEK bulk chromatin *(top panel)* and H3.3-containing nucleosomes from HEK *(bottom panel)*. Yellow box identifies H2A.Z variant detected at 14% abundance in H3.3-containing nucleosomes. All proteoform peaks are normalized to the intensity of the peak of H2A.A/O (n = 3). **(g)** Example tracks from ChIP-seq reads in HEK cells showing input, H3.3 and H2A.Z targets supporting co-localization of these two variants. **(h)** Heatmap centered on H3.3 peaks ±0.5 kb showing the correlation of ChIP-seq signal between H3.3 and H2A.Z (antibodies from Abcam and Cell Signaling Technologies, CST). Clusters of loci are compositionally defined in **Supplemental Fig. 10** and described with gene ontology terms in **Supplemental Fig. 19**.

To test the ability of Nuc-MS to localize PTMs, this three-stage process was used to measure the effects of histone-modifying enzymes on nucleosome substrates. We first interrogated nucleosomes acetylated by p300/CBP associated Factor (PCAF), which plays key roles in transcription and DNA repair.^14^ A synthetic unmodified nucleosome was assembled and then treated with the catalytic domain of PCAF/KAT2B (aa 492-658) for five minutes at room temperature. Nuc-MS analysis of PCAF-treated nucleosomes revealed >95% of monoacetylation to be at H3K14ac, consistent with the known substrate specificity of the enzyme (**Supplemental Fig. 5**).^14^

We also used Nuc-MS to delineate the products of the Polycomb repressive complex 2 (PRC2) after incubation with synthetic nucleosomes for 18 hours (**Supplemental Fig. 6**). PRC2 is the only enzyme responsible for H3K27 methylation in humans,^15^ and plays a central role in transcriptional repression and cellular differentiation, with its dysregulation linked to several types of cancer.^16^ Nuc-MS analysis showed >95% of the methylation was in fact limited to H3K27, with the reaction products created by the action of the EZH2-SET domain being 40% H3K27me1, 50% H3K27me2 and 2% H3K27me3 (**Supplemental Fig. 6**). Combining Nuc-MS with nucleosomes defined by PTMs at specific amino acid sites^18^ will enable nucleosome-level research in the fields of epigenetics^19^ and proteomics.^5^ Nuc-MS will be complementary to conventional assays that use short histone peptides to investigate PTM reading, writing, and erasing, as these experiments do not yet account for the whole nucleosome context and chromatin landscape.^19, 20^

Given the relative simplicity and low bias of Nuc-MS to capture the proteoform landscape of dominant histone proteoforms in a population of nucleosomes, we challenged the approach with all the variants and PTM patterns present on endogenous nucleosomes from human HEK 293T and HeLa cells (**Fig. 2** and **Supplemental Fig. 7**). The MS^1^ measurement of intact mono-nucleosomes produced by Micrococcal nuclease (MNase) treatment resulted in a spectrum with no visible charge states for mass deconvolution (**Supplemental Fig. 8a**). However, using the new approach of Individual Ion MS (I^2^MS)^21^ we were able to precisely profile the mass distribution of these mononucleosomes, centered at ∼200 kDa (**Supplemental Fig. 8b-d**).

Upon activation of the *entire* population of mononucleosomes for the tandem MS experiment, it proved straightforward to eject and detect >70 histone proteoforms at high resolution in the same spectrum (**Fig. 2a**). Examination of spectral regions for each core histone reveals a clear snapshot of all proteoform distributions present in bulk chromatin at >1% relative abundance (**Fig. 2b-d**).^5^ Importantly, the quantitative readout of H2A and H2B distributions of gene family members achieved by Nuc-MS would be challenging to obtain by other proteomic techniques. Additionally, the major proteoform of histone H4 was characterized by tandem MS^3^ as H4K20me2-N_t_ac (**Fig. 2e**, **Supplemental Fig. 7e**). By simultaneously detecting the dominant histone proteoforms, this new data type provides a low-bias, isoform-specific, and quantitative survey of the epigenetic landscape in global chromatin without upfront chromatography or data ‘recombineering’. The method is broadly applicable to mononucleosomes obtained from diverse tissues, model organisms or affinity-enrichment.

We next used Nuc-MS to report on selected types of nucleosomes present in sub-regions of chromatin. To this end, we transfected HEK cells with a plasmid containing the *H3.3F3A* gene with a C-terminal extension encoding a FLAG-HA tag. We then prepared nucleosomes and immuno-enriched for H3.3-FLAG, which is known to localize in euchromatic regions.^22^ Upon comparison of the histone profiles from FLAG-enriched nucleosomes and from HEK bulk chromatin as a control (**Fig. 2f, Supplemental Fig. 9**), we identified major differences, including ∼6-fold enrichment of H2A.Z in H3.3 nucleosomes (*p* = 2.7×10^-7^), and elevation of unmodified H4, H4K20me1, and acetylated H4K20me1 proteoforms in H3.3 nucleosomes by 80 ±3.4%, 40 ±0.7%, and 37.5 ±1.2%, respectively (with *p* = 7.5×10^-6^, 2.3×10^-6^, and 6.8×10^-5^). Nuc-MS is the first technique to date that is able to directly quantify the extent of H3.3 and H2A.Z co-enrichment; indeed, ∼1 out of 7 nucleosomes was found to contain both H3.3-FLAG and H2A.Z (**Fig. 2f**, **yellow box at left**). In the same experiment, we detected co-occurring proteoforms that were elevated above their levels in bulk chromatin and that have been highly correlated with regions of active transcription and high H3.3-H4 tetramer turnover.^23–25^ To validate our findings, we analyzed H3.3 and H2A.Z by ChIP-seq.^22^ Representative loci show highly similar tracks for H3.3 and H2A.Z peaks (**Fig. 2g**). The extent of co-occupancy for these two variants is captured by the heatmap which aggregates ∼73,000 reads into four clusters (**Fig. 2h**). The elevated signal intensity in clusters 1-3 reflects up to 20% co-occurrence for H3.3 and H2A.Z in introns and promoters (**Supplemental Fig. 10**); this is consistent with the Nuc-MS measurement of 13.7 ±0.2% co-occurrence (**Fig. 2f**).

Finally, Nuc-MS was used to profile the composition of nucleosomes harboring H3.3 K27M. This ‘toxic’ oncohistone arises in >80% of diffuse intrinsic pontine gliomas (DIPG), a highly aggressive tumor of the pediatric brain stem.^26, 27^ Previous work found that H3.3K27M associates with K27ac on H3WT, and that the mutation and K27ac co-localize with RNA pol II, indicating that these marks are present in sites of active transcription.^27, 28^

As above, we performed immuno-precipitation of mononucleosomes from HEK 293T cells expressing inducible transgenes H3.3 K27M-FLAG-HA and the control H3.3 WT-FLAG-HA (**Supplemental Fig. 11**). Selective activation of the K27M and WT mononucleosomes enabled the wide-lens profiling of their histone proteoforms (**Fig. 3a, 3b**). Quantitative analysis of the relative proteoform abundances revealed significant differences between K27M and WT nucleosomes. Importantly, H3.3K27M nucleosomes contain a 33.7 ±1.4% increase in H4K16ac-N_t_ac, a proteoform correlated with active transcription and increased accessibility of H4 to acetyl-transferases^29^ (*p* = 1.01×10^-4^; **Supplemental Fig. 12**; α-values reported in **Supplemental Tables 1-2**).

**Figure 3.**
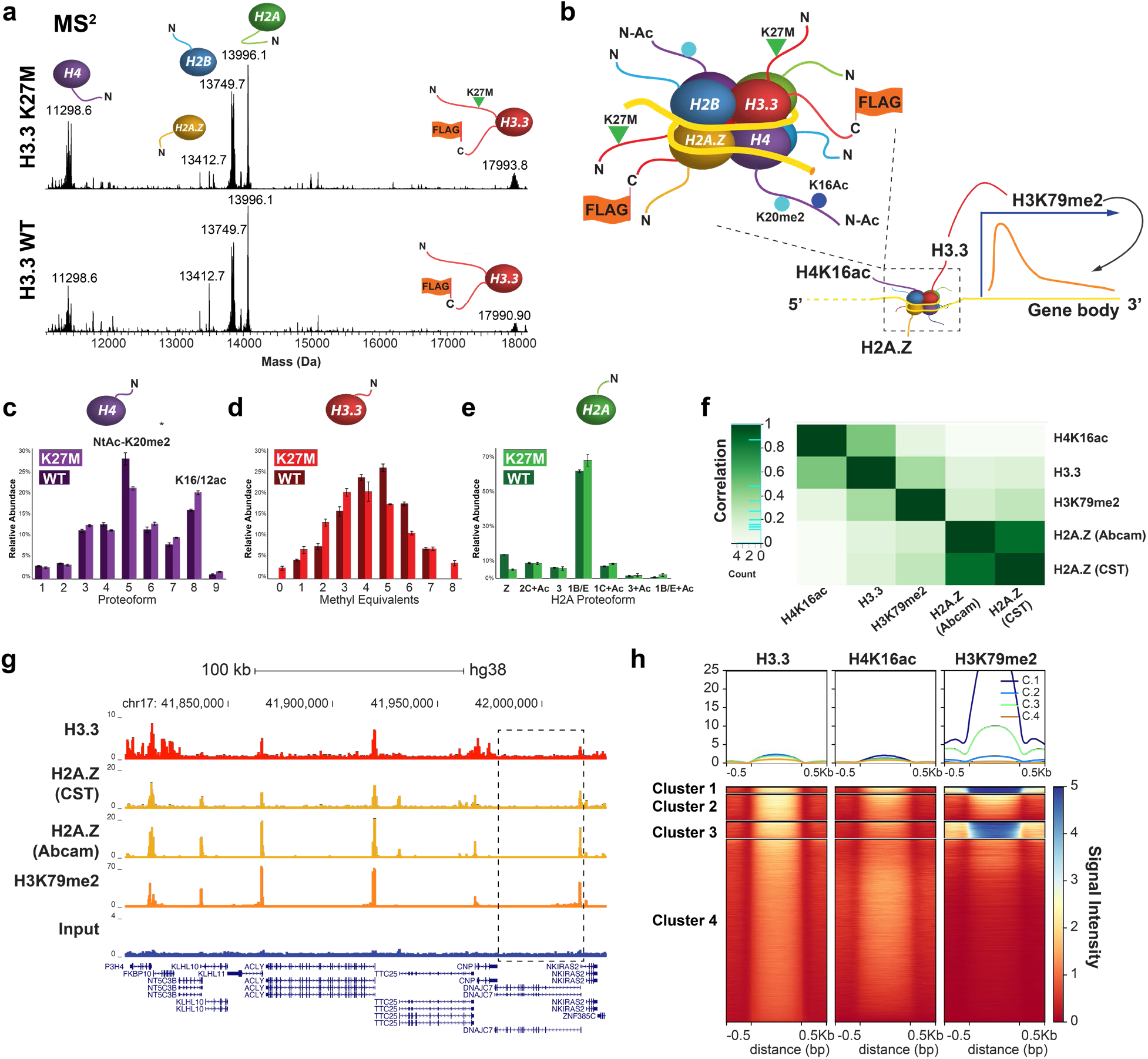
Nuc-MS of endogenous nucleosomes prepared from cells with WT or K27M forms of histone H3.3. **(a)** MS^2^: measurement of the histone proteoforms ejected from mononucleosomes isolated from two cell lines (all measured at isotopic resolution; 2 biological replicates and 3 measurement replicates). Mononucleosomes were isolated from 6000-9000 *m/z* to eject histone proteoforms for MS^2^ measurement. Note the ∼2.5 kDa shift in H3 due to the addition of the FLAG-HA-tag in comparing the MS^2^ spectra of H3.3-enriched HEK vs HEK bulk mononucleosomes in Figure 2. (**b)** Depiction of the composition for the most abundant nucleosomes determined by Nuc-MS, reflecting high enrichment for histone proteoforms and variants present at promoters and highly expressed genes. **(c-e)** Quantitative analysis of proteoform abundances for ejected histone proteoforms by MS^2^, revealing the co-occurrence of H3.3K27M with proteoforms and PTMs consistent with transcriptional activation. In panel **(d),** the WT data for H3.3 was shifted by +3 Da (mass of K→M mutation) in order to better illustrate proteoform differences when comparing with H3.3K27M. Asterisks in panel (d) denote artifacts from the charge deconvolution process. **(f)** Pearson correlation plot showing association among histone marks and variants characterized by Nuc-MS and targeted by ChIP-seq (H3.3, H3K79me2 and H2A.Z, and H4K16ac). **(g)** Example tracks showing ChIP-seq reads in HEK cells for input, H3.3, H3K79me2 and H2A.Z support their co-localization. A zoom-in of the gene highlighted with a black box is provided in **Supplemental Fig. 18**. **(h)** Heatmap centered on H3.3 peaks ±0.5 kb showing the correlation of ChIP-seq signal between H3.3, H3K79me2 and H4K16ac. A complete heatmap and additional cluster information is provided in **Supplemental Fig. 10**, with gene ontology and relevant biological pathways listed for each cluster in **Supplemental Fig. 19**.

The data further reveal a hypo-methylated state of H3.3K27M in relation to WT H3.3 that results from the loss of the K27 methylation site (**Fig. 3d**). Specifically, the average number of methyl equivalents decreases from 4.4 ±0.02 in H3.3WT to 3.9 ±0.1 in H3.3K27M nucleosomes (n = 3). In terms of nucleosome symmetry, we detected far lower levels of WT H3 from H3.3K27M-FLAG and H3.3WT-FLAG, as confirmed by SDS-PAGE, indicating that these nucleosomes are >90% homotypic for H3.3-FLAG tails (**Supplemental Fig. 13**). Given the low expression levels of H3.3-FLAG constructs relative to total H3 upon doxycycline induction,^27^ the predominance of homotypic (2x) FLAG-tagged nucleosomes may have resulted from avidity bias during immunoprecipitation. Though this biochemical limitation prevented detection of K27ac or K27me3 marks on the endogenous H3WT tails, the data validate that Nuc-MS can quantitatively interrogate the homotypic vs. heterotypic nature of nucleosomes.

Fragmentation of the H3.3K27M-FLAG proteoforms enables characterization of methylation. For example, five informative ions shown in **Supplemental Fig. 14** reflect the very high co-occurrence of K27M with H3K79me2, a mark correlated with active transcription. This finding is notable as H3.3K79me2 has been found to be <2.5% of H3.3K79 methylation,^30^ and is therefore enriched in H3.3K27M nucleosomes by >15-fold relative to bulk (**Supplemental Fig. 14**). Additionally, fragmentation of H3.3K27M indicates that the most abundant proteoform also contains large amounts of two to three methyl equivalents in the region between residues 23-72; this is most likely an elevation of K36me2/3, which has been detected previously with the K27M mutation but only asserted to co-occur in the same nucleosomes by extensive use of ChIP-seq.^28^ Supporting the Nuc-MS data, analysis of the tryptic peptides of histones from HCT116 cells reveals elevation of both H3.3K79me2 and H3.3K36me2/3 (**Supplemental Figure 15**).

Examination of H2A proteoforms revealed a modest elevation in H2A acetylation and nearly a three-fold decrease in H2A.Z abundance (**Fig. 3e**) in H3.3K27M-FLAG nucleosomes compared to H3.3WT-FLAG (5.1 ±0.2% vs. 13.7 ±0.03%, n = 3; *p* = 1.7×10^-4^; **Supplemental Fig. 16**). Moreover, H2A.Z acetylation, which is 3.3 ±0.4% (n = 3) of total H2A.Z in H3.3WT, decreases to undetectable levels in K27M nucleosomes. This result suggests that K27M may be correlated with H2A.Z eviction from the nucleosome, which presents functional implications.^31, 32^ Finally, we did not detect ubiquitinated histones in H3.3WT or K27M nucleosomes despite the potential for cross-talk^33^ (**Supplemental Fig. 17**).

We again turned to ChIP-seq to validate our Nuc-MS results and interrogate the extent of co-localization of H3.3 variant with H4K16ac, H3K79me2, and H2A.Z. The ChIP-seq data are summarized by a Pearson correlation analysis (**Fig. 3f**), example tracks from selected loci (**Fig. 3g**), and a heatmap centered around H3.3 (**Fig. 3h**), that together show strong co-association and inform a model for the composition of nucleosomes at promoters (**Fig. 3b**). H4K16ac shows the strongest Pearson correlation to H3.3 (0.48, **Fig. 3f**); this is consistent with the heatmap profiles for both marks, which show signal intensity across the whole genome and no cluster-specific enrichment (**Fig. 3h**). In contrast, the heatmap reveals that H3K79me2 signal is mostly localized to cluster 2, made up of 45% promoters and 22% introns (**Supplemental Figs. 10, 18, 19**). Close inspection of a representative gene shows that H3K79me2 displays a prominent promoter-associated peak that co-localizes with H3.3 and H2A.Z;^34^ however, unlike these variants, H3K79me2 signal extends beyond the transcription start site into the gene body (**Supplemental Fig. 18**).

In sum, H3.3K27M nucleosomes show a clear co-occurrence with histone proteoforms and PTMs consistent with a euchromatic state that have not been previously reported. The nucleosomal characteristics that follow the loss of the H3K27 methylation site (i.e. lower methylation state of H3, >15-fold enrichment of H3K79me2 relative to bulk, increased acetylation of H4 and H2A), is consistent with a model of active transcription and chromatin de-condensation.^29^ Nuc-MS showed high agreement with orthogonal readout by ChIP-seq, and constitutes a direct measurement of histone variants, proteoforms and their PTMs that co-occur in whole nucleosomes. Finally, combining this approach with immuno-enrichment of rare marks will help develop new insights into the role of a ‘nucleosome code’, where co-occurring histone proteoforms combine to potentiate regulation of gene expression and disease phenotypes.

## Materials and Methods

### Nucleosome Assembly

Nucleosome particles were assembled by salt dialysis.^1^ Specifically, resuspended 50 µg of 601 nucleosome positioning sequence DNA (*EpiCypher*) in 50 µL nuclease-free 2M NaCl. Heated to 37 °C and mixed thoroughly. Then mixed the following reagents, in order: 54.2 µL 2M NaCl, 50 pmol 601 DNA (45.8 µL), 50 µL of 20 µM H2A/H2B dimer (*New England BioLabs*), and 50 µL of 10 µM H3/H4 tetramer (*New England BioLabs*). The NaCl concentration was gradually lowered by adding increasing volumes of 10 mM Tris HCl (pH 8) every thirty minutes (final concentrations at each step: 2M; 1.48M; 1.0M; 0.6M; 0.25M). Sample was then added to a 20 kDa MWCO Slide-A-Lyzer MINI dialysis device (*Thermo Fisher Scientific*) and dialyzed against three buffer changes (the second one being overnight) of 100 mM ammonium acetate (pH 6.8). After dialysis, samples were exchanged into 10 mM Tris HCl, pH 8 and concentrated to 100 µL using 30 kDa MWCO spin filters (Millipore-Sigma).

For MNase digestion, samples were brought to 200 µL with 10 mM Tris buffer (pH 8), and supplemented with 4 µL of 100 mM CaCl_2_ (2 mM final), and 20 µL of 20 U/µL MNase (*New England BioLabs*, 1/1000 dilution of commercial stock). Reactions were digested for 1 minute and quenched with 2 µL 500 mM EDTA and thorough pipetting; and desalted 4-5 times into 150 mM ammonium acetate (pH 6.8) using 100 kDa MWCO spin filter. Effective nucleosome assembly was confirmed by native TBE gels (*BioRad*).

The nucleosome ubiquitinated at H2AK119 (*EpiCypher* #16-0363) and mono- and di-methylated at K36 (*EpiCypher* #16-0322 and #16-0319) were desalted 10 times into 150 mM ammonium acetate using 30 kDa MWCO spin filters prior to MS analysis.

### Nucleosome Enzymatic Modification (PCAF and PRC2)

#### PCAF

For nucleosome acetylation, the following reaction was set up: 50 µL 5x Histone Acetyltransferase (HAT) buffer (250 mM Tris HCl, 50% glycerol, 0.5 mM EDTA, 5mM DTT), 8 µL Acetyl CoA at 10 mM, 40 µL of 2.8 µM nucleosome, 2 µL recombinant PCAF (aa 492-658) at 197 µM, 150 µL 150 mM ammonium acetate. Mixture incubated at RT for 5 minutes and desalted using 30 kDa MWCO spin filters (*Millipore-Sigma*) into 150 mM ammonium acetate for MS analysis.

#### PRC2

for nucleosome methylation, the following reaction was set up: 50 µL nucleosome at 2.8 µM, 1 µL 100 mM TCEP, 0.2 µL 1M MgCl_2_, 3.33 µL 1.5 M Tris (pH 8.8), 1 µL 10 mM SAM, 2 µL PRC2 (2.8 µM, Active Motif), 42.47 µL water. Mixture was incubated at RT for 18 hours and desalted into 150 mM ammonium acetate for MS analysis using 100 kDa MWCO spin filters (*Millipore-Sigma*).

### Preparation of HEK and HeLa Mononucleosomes

Harvested cells were resuspended in 2.5 pelleted cell volumes (PCV) of buffer A (10mM HEPES, pH 7.9, 10mM KCl, 340mM Sucrose, 0.5mM PMSF, 0.5mM Benzamidine) supplemented with 5 mM βME and 1x cOmplete^TM^ EDTA-free protease inhibitor cocktail (Roche). A similarly supplemented buffer A volume (2.5 PCV) containing 0.2% (v/v) Triton X-100 detergent was also prepared during this time. The resuspended cell pellet was completely homogenized by pipetting and light vortexing and the detergent containing buffer A was added to the cell suspension, mixed with a few inversions, and allowed to incubate on ice for 10 minutes with occasional gentle mixing to thoroughly lyse the cells. The resultant nuclei were pelleted via centrifugation (1300 x g, 4 °C, 5 minutes) and the supernatant was removed via aspirartion and discarded.

The nuclei pellet was next resuspended in 6 PCV of supplemented buffer A and spun through a 35 mL sucrose cushion (10 mM HEPES pH 7.9, 30% (w/v) sucrose) in a 50 mL conical tube supplemented with 5 mM βME and 1x cOmplete^TM^ EDTA-free protease inhibitor cocktail (1300 x g, 4 °C, 10 minutes). The sucrose cushion purification of the nuclei was repeated until the cushion remained free of cell debris after centrifugation.

The pelleted nuclei were then gently resuspended in 2 PCV of buffer A (supplemented as above). The DNA concentration in the nuclei resuspension was calculated by hypertonic lysing of the nuclei by mixing 5 µL of the nuclei resuspension with 495 µL 2M NaCl. DNA concentration was determined via A260 measurements. The suspension was diluted to achieve a DNA concentration of 1.5 μg/μL and CaCl_2_ was added to the nuclei to a final concentration of 1 mM. The suspension was aliquotted into 2 mL microcentrifuge tubes, and equilibrated in a 37 °C water bath for 5 minutes. Digestion of the nuclei to nucleosomal species was initiated by adding 1U MNase (Worthington; prepared in supplemented buffer A) for every 70 μg of DNA and the samples were allowed to incubate at 37 °C for 15 minutes. After the incubation period, the digestion was quenched by adding 0.5 M EGTA solution to a final concentration of 10 mM, mixing via inversion, and placing the samples on ice. The final volume of the digested material was noted. Approximately 10 µg of DNA was purified via QIAGEN DNA cleanup kit and resolved on a 2% agarose gel in 0.5x TBE to confirm extent of digestion.

2M NaCl was added dropwise to the digested chromatin while mixing on a magnetic stir plate at 4 °C to a final concentration of 650 mM. The resulting material was cleared via centrifugation at 12,000 x g at 4 °C to pellet any insoluble material before size exclusion chromatography. The material was fractionated via a HiPrep™ 26/60 Sephacryl® S-300 HR column equilibrated with supplemented buffer A, as above, containing 650mM NaCl using an ÄKTA Prime Plus FPLC (GE Lifescience). 10 µg samples of the individual fractions containing nucleosomes were purified via QIAGEN DNA cleanup kit and resolved on a 2% agarose gel in 0.5x TBE. Only those fractions that contained mononucleosomal associated DNA fragments (∼150bp) were pooled and used for further analysis.

### Mononucleosome Flag-immunoprecipitation

Immunoprecipitation of Flag-tagged H3.3WT and H3.3K27M histones was performed as previously.^27, 28^

### Chromatin Immunoprecipitation for ChIP-seq

HEK293T cells (1×10^8^) were crosslinked with 1% formaldehyde for 15 minutes and quenched with 0.125M glycine. Cell lysis and chromatin preparation were performed as previously described (Lee et al., 2006; Vo et al., 2017). For sonication, the sample was resuspended in 2ml of ChIP buffer #3 and sonicated on a Covaris E220 sonicator for 5 minutes, 200 cycles per burst, 140W peak intensity pulse and 20% duty factor. Following sonication, samples were cleared by centrifugation at 20,000xg for 15 minutes. Chromatin was then diluted to a concentration of 0.4mg/ml and Triton X-100 was added to a final concentration of 1%. For each ChIP binding reaction, 0.4mg of chromatin was incubated with 10ul of antibody overnight at 4C. The following day, 20ul of Protein-A/G plus beads (Santa Cruz sc-2003) were added to each ChIP and incubated for 2 hours. Beads were washed 5 times in ChIP wash buffer and 2 times in TEN buffer. ChIPed DNA was purified using phenol: chloroform as described (Lee et al., 2006; Vo et al., 2017). Full references available in **Supplemental Information**. Antibodies used: H2A.Z (Cell Signaling Technologies, CST, #2718S) Lot #2; H2A.Z (Abcam, #ab4174) Lot# GR3198864-2; H3.3 (EMD/Millipore, #09-838) Lot #3310680; H4K16ac (CST, #13534S) Lot #3; H3K79me2 (CST, #5427S) Lot#4.

### ChIP-sequencing and data processing

ChIP-sequencing libraries were synthesized using the Illumina TruSeq kit, size selected (200-400bp) with SPRI select beads and sequenced on an Illumina Novaseq instrument. Base-calling was performed using bcl2fastq and read quality was assessed with FastQC (Andrews, 2010). ChIP-seq reads were aligned to the human genome (hg38) using Bowtie v0.12.9 (Langmead et al., 2009) allowing for 2 mismatches and retaining only uniquely mapped reads. MACS v2.1.0 (Zhang et al., 2008) was used to call peaks using a false discovery rate filter of 0.01. Annotation of ChIP-seq data was performed using HOMER v4.10 (Heinz, 2010) and Pearson correlation for samples was calculated using R package DiffBind (Ross-Innes et al. 2012). Functional gene enrichment analysis was performed using R package clusterProfiler (Yu et al., 2012). Genome wide occupancy heatmaps were generated using deepTools v3.1.1 (Ramirez et al., 2016) and centered on H3.3 peaks. Full references available in the **Supplemental Information**.

### Native Mass Spectrometry

Samples were analyzed using a Q Exactive HF mass spectrometer with Extended Mass Range (QE-EMR) and a Q Exactive HF Ultra-High Mass Range (QE-UHMR), both by *Thermo Fisher Scientific*. Data were collected using XCalibur QualBrowser 4.0.27.10 (Thermo Fisher Scientific). The native electrospray platform is coupled to a three-tiered tandem MS process. First, the analysis of the intact nucleosome (MS^1^) provides the total complex mass (reported as a deconvoluted neutral average mass value) ^11^. In stage two, the nucleosome is activated by collisions with nitrogen gas to eject histones (MS^2^). In stage three, further vibrational activation of the ejected histones via collisions with nitrogen gas yields backbone fragmentation products from each monomer (MS^3^) that are recorded at isotopic resolution (120,000 resolving power at *m/z* 400). These fragments can be mapped onto the primary sequence of the histones in order to localize posttranslational modifications.

#### QE-EMR parameters

The Nuc-MS workflow utilizes native electrospray ionization (nESI) source held at +2 kV, C-trap entrance lens voltage setting between 1.8 - 4 V, HCD gas pressure setting between 2-4 V, and CID voltage set at 15-25 V for desalting and 75-100 V for histone ejection. HCD energy set to 100-120 V for histone fragmentation with a pressure of 2. Microscans set to 20 and max injection time to 2000 ms for collection of fragmentation data.

#### QE-UHMR parameters

HCD gas pressure between 0.5-1 for detection of histones at isotopic resolution; in-source trapping voltage of −100 to −150 V for histone ejection; CE of 49-65 eV, microscans set to 20 and max inject time to 1000 ms for fragmentation of proteoforms.

Prior to using the QE-UHMR for nucleosome analysis, quantitative ejection of the PCAF- and PRC2-treated histones for fragmentation was achieved using front-end infrared activation coupled to the QE-EMR, which used a 20 W continuous-wave CO_2_ laser (*Synrad Firestar V20*) at an average power of 1.2 W. The laser was attenuated with a 1.0 optical density (O.D.) nickel-coated zinc selenide neutral density filter and aligned unfocused to the inlet capillary with protected gold mirrors. Once we transitioned the Nuc-MS platform to the QE-UHMR, the in-source trapping capability proved to be a reliable method for ejecting histones at sufficient intensities for adequate fragmentation.

### MS Data Analysis

Intact mass values for nucleosome complexes and ejected histones, the MS^1^ and MS^2^ measurements, were determined by deconvolution to convert data from the *m/z* to the mass domain using MagTran 1.03^35^ (mass range: 15,000-300,000 Da; max no. of species: 10-15; S/N threshold: 1; mass accuracy: 0.05 Da; charge determined by: charge envelop only). Intact mass measurements are reported as neutral average masses; errors represent 1σ deviation from the mean of the masses calculated for of all sampled charge states.

UniDec 3.2.0^36^ was used to create isotopically resolved deconvoluted mass spectra, depicted in the butterfly diagrams. Data processing: Range 500 – 2500 Th, Bin every: 0; UniDec parameters: Charge Range: 5 – 15, Mass Range: 10 – 20k Da, Sample mass every (Da): 0.05. Peak area values were calculated by integrating the assigned mass range of proteoforms in the deconvoluted mass spectrum. Bar diagrams for relative quantification of proteoforms were based on direct infusion experiments (n = 3) and mean relative peak area values with standard deviations as error bars. Statistical significance was evaluated using two-sided, two-sample t-tests. The α-values (based on α = 0.05) were adjusted according to the Bonferroni correction and both α- and *p*-values are reported in **Supplemental Tables 1-2**.^37^

High-resolution fragmentation data were processed using Xtract (Signal-to-Noise threshold ranging from 1-30, *Thermo Fisher Scientific*), mMass 5.5.0 (www.mmass.org), ProSight Lite 1.4^38^ (precursor mass type: average; fragmentation method: HCD; fragmentation tolerance: 10-15 ppm), and TDValidator 1.0^39^ (max ppm tolerance: 25 ppm; cluster tolerance: 0.35; charge range: 1-10; minimum score: 0.5; S/N cutoff: 3; Mercury7 Limit: 0.0001; minimum size: 2) to assign recorded fragment ions to the primary sequence of the subunits. Specifically, ProSight Lite and TDValidator were used to analyze fragmentation spectra in medium throughput to assign and validate *b* and *y* fragment ions to the histone sequences, and for generating a p-score. mMass was used to interrogate individual fragment ions within a spectrum not identified by TDValidator or ProSight Lite. The histones H2A, H2B, H3, and H4 were identified by mapping backbone fragment ions to their amino acid sequence using ProSight Lite.^38^ Unexplained mass shifts (Δ*m*) observed at the MS^1^, MS^2^, and MS^3^ levels for the intact complex and subunits, respectively, were manually interrogated using the UNIMOD database (http://www.unimod.org/modifications_list.php) as a reference for candidate modifications

### MS Data availability

MS^1^, MS^2^, and MS^3^ spectra presented in the manuscript will be made available online in the MassIVE database under accession code MSV000085238 after acceptance of this work in a peer-reviewed journal.

### Ion Collection and Data Acquisition for Individual Ion Mass Spectrometry (I^2^MS)

Detailed methods for this new technique are reported in Kafader, *et al.* (2019).^21^ Briefly, this new method uses direct assignment of charge states on individual ions inside an Orbitrap-style mass spectrometer with a harmonic potential. To provide populations of single ions of endogenous mononucleosomes, the transmission of the instrument was detuned to lower the number of ions entering the Orbitrap analyzer and achieve detection of a single ion per *m/z* value for each acquisition event (disabled Automatic Gain Control and maximum injection time between 0.03 - 1 ms). Both the *m/z* and charge (*z*) were necessary to determine the mass of the ion. In the Orbitrap portion of a Q Exactive instrument (*Thermo Fisher Scientific*), the *m/z* of ions were determined from the frequency of ion rotation around the central electrode; the charge, *z*, was given by the rate of the induced charge on the outer electrode, also known as Selective Temporal Overview of Resonant Ions (STORI), described in detail elsewhere.^21^ Plotting of the I^2^MS spectrum from this multiplexed, I^2^MS procedure was achieved by binning ∼1M acquired individual ions of mononucleosomes in 0.2 Da increments. In parallel, to validate the charge assignment, the calculated charge of the ions used for the individual ion MS spectrum were binned in quantized domains, as reported previously (data not shown).

## Acknowledgments

This work was supported by the National Institute of General Medical Sciences P41 GM108569 for the National Resource for Translational and Developmental Proteomics at Northwestern University and NIH grants S10OD025194 and RF1AG063903 (Kelleher lab) and R44GM116584, R44CA212733 and R44CA214076 (EpiCypher). LFS is a Gilliam Fellow of the Howard Hughes Medical Institute. Research in this publication is also supported by Thermo Fisher Scientific and a fellowship associated with the Chemistry of Life Processes Predoctoral Training Grant T32GM105538 at Northwestern University. A.P. is supported by the transition to independence grant K99CA234434-01. Bottom-Up proteomics services were performed by the Northwestern Proteomics Core Facility, generously supported by NCI CCSG P30 CA060553 awarded to the Robert H Lurie Comprehensive Cancer Center. We also thank Michael Senko, Philip Compton, Christopher W. Koo, Lindsey C. Szymczak, and Michael O. McAnnally for technical assistance, and Sheila Judge and Amy Rosenzweig for providing thoughtful suggestions to the manuscript.

## Author Contributions

Data acquisition and analysis was performed by LFS. KJ assisted with proteoform quantitation and MS analysis. AP, MM, AL and AS prepared and made available H3.3K27M and WT mononucleosomes. JOK assisted with acquisition of multiplexed single-ion MS data on endogenous nucleosomes. MJM, MAC, JMB and SAH assisted with synthesis, purification, and verification of modified nucleosomes. MCK coordinated modified nucleosome synthesis and provided insightful feedback on the manuscript. LFS and NLK conceived of the project and wrote the manuscript.

## Supplemental Information

**Supplemental Figure 1.**
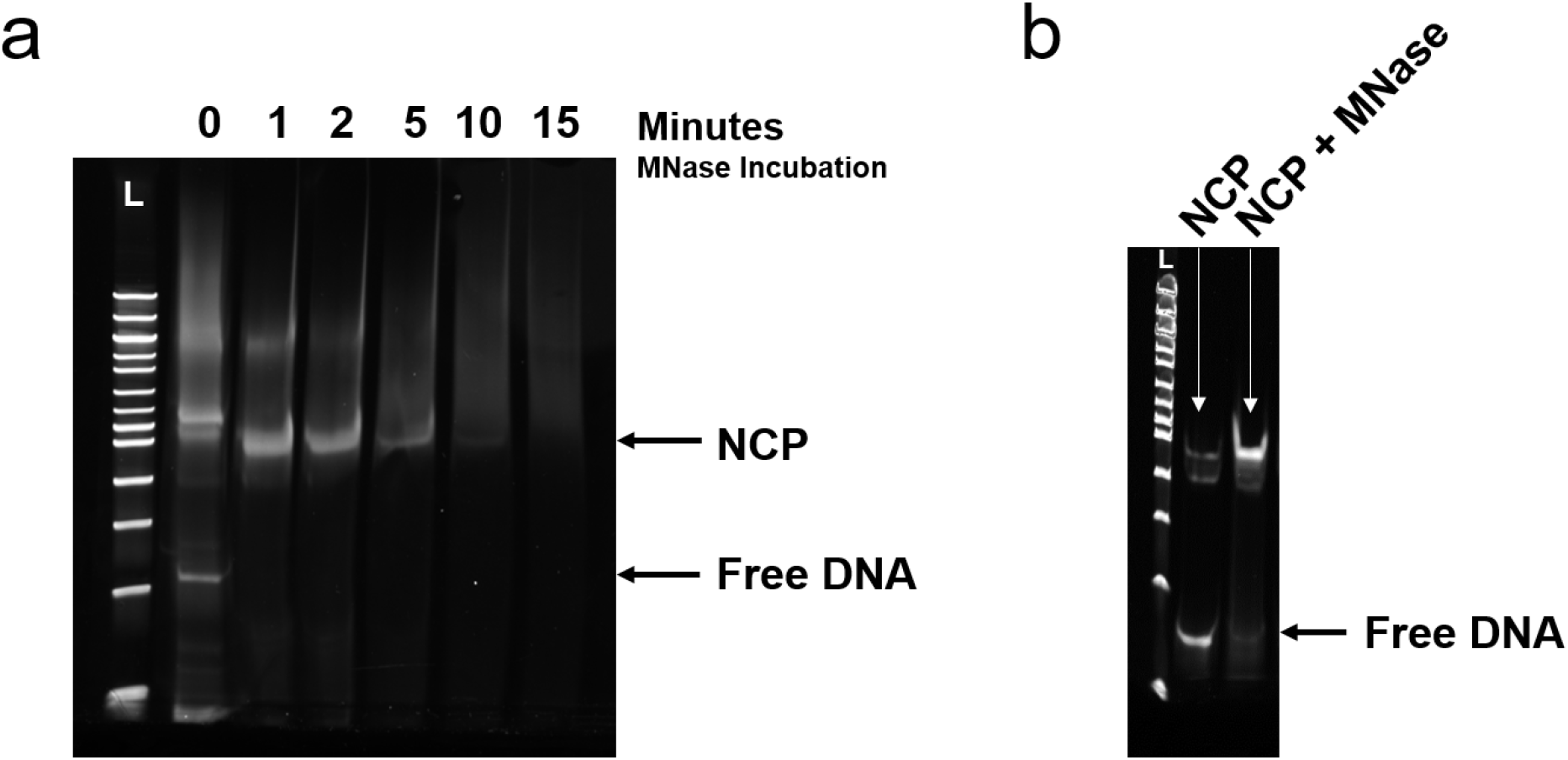
Synthetic nucleosomes were assembled by salt dialysis and MNase treated to eliminate free DNA prior to analysis by Nuc-MS (employing native electrospray).^40^ **(a)** Nucleosome core particle (NCP) samples with 208 bp DNA were treated with 20U of MNase for 0-15 minutes, and quenched with 25 mM EDTA. Samples are resolved by TBE native PAGE and stained with GelRed (*Biotium*). **(b)** NCP with 147 bp DNA treated with MNase for 1 minute. The bp ladder lane is indicated by the letter “L”.

**Supplemental Figure 2.**
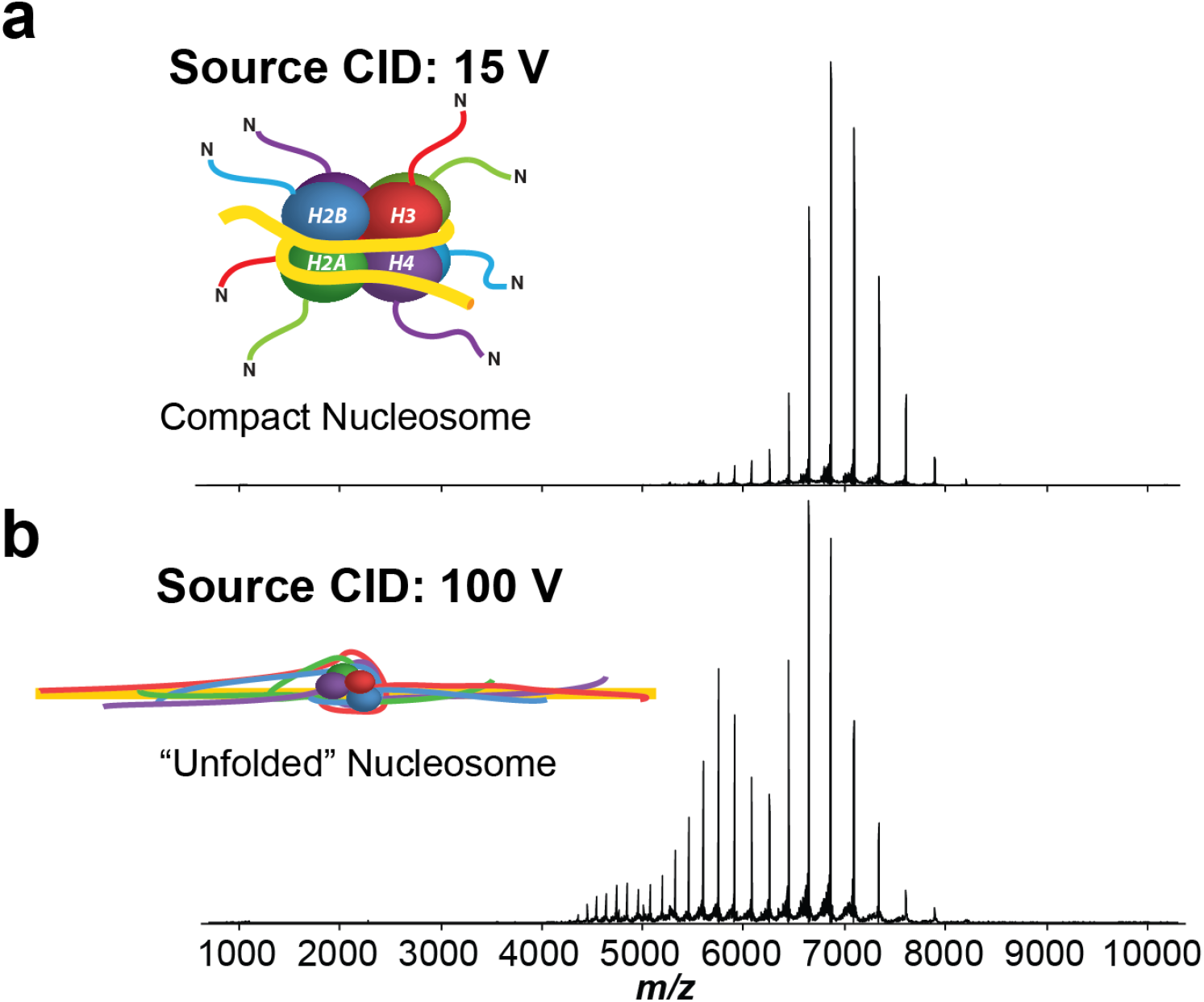
“Unfolding” of the nucleosome in the Electrospray source. MS^1^ of nucleosome at low **(a, 15 Volt)** and high **(b, 100 Volt)** source activation energy (source-induced dissociation, source CID). An increase in collisional energy results in a concomitant increase in the average charge state and width of the charge state distribution in the native mass spectrum; spray solution was pH ∼7, 150 mM ammonium acetate. This observation is consistent with a current model of nucleosome unfolding,^41^ where the DNA in the nucleosome core particle first linearizes upon activation, thereby extending the strongly-bound histones across the length of the strand, and potentially increasing exposure of additional sites to protonation during electrospray ionization.

**Supplemental Figure 3.**
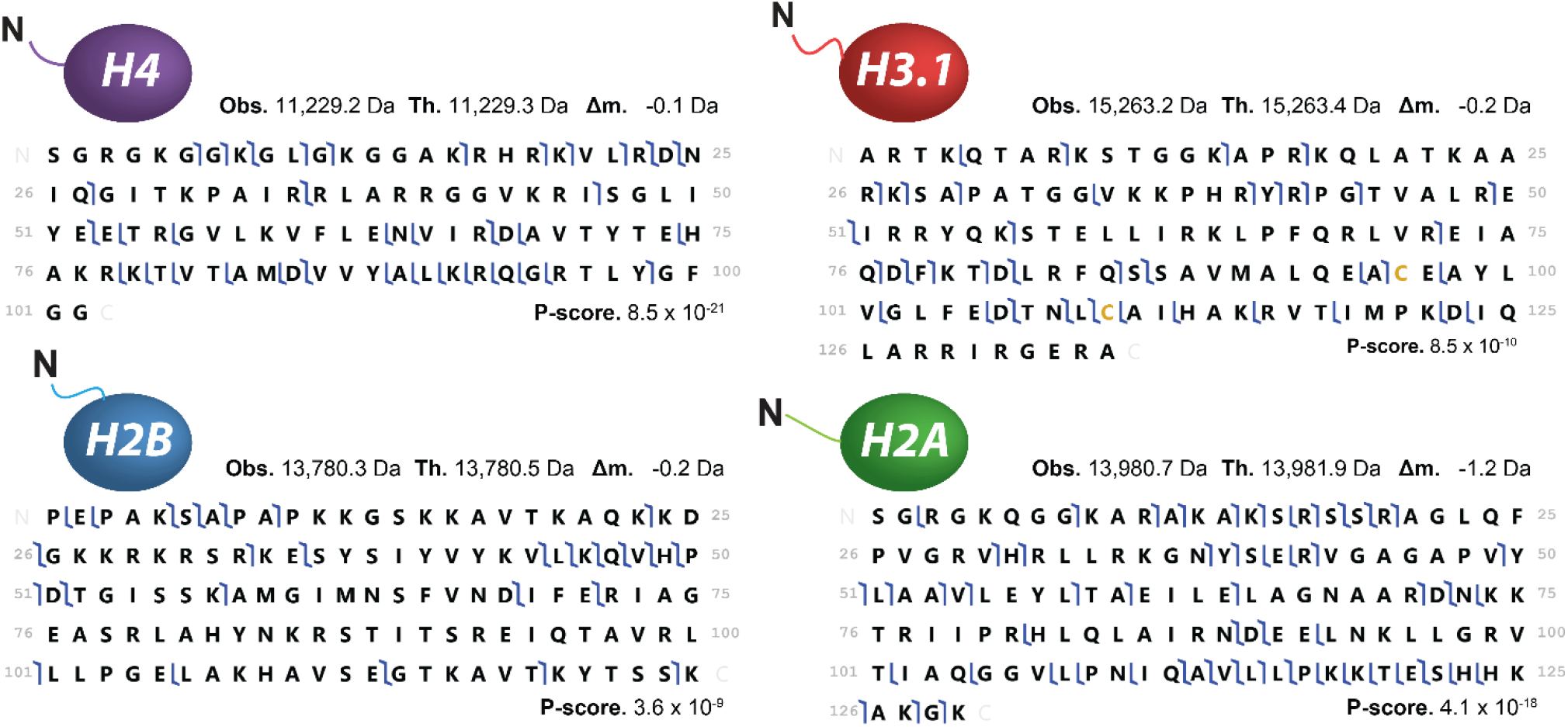
Fragment maps for all four histones ejected from unmodified recombinant nucleosomes (**Figure 1b**). Fragments were manually validated using TDValidator.^42^ P-scores were calculated using ProSight Lite.^43^

*Supplemental Discussion for Nuc-MS of unmodified recombinant nucleosomes (Figure 1 and Supplemental Figures 2-3).* After measuring the intact nucleosome mass(es), and then liberating core histones and their sequence ions, Nuc-MS provides isoform and variant-specific identification, with PTM localization (i.e. the complete proteoform; **Fig. 1b**, and **Supplemental Figs. 1** and **2**). The theoretical average mass of a recombinant nucleosome, comprising eight histones (108,576.6 Da) and 147 bp of biotinylated 601 DNA nucleosome positioning sequence (91,294.5 Da), was calculated to be 199,871.1 Da.^40^ Deconvolution of the charge states in the MS1 spectrum yielded an observed average mass of 199,867.9 ±12.5 Da, a −3.1 Da difference (Δm) from the theoretical value (**Fig. 1b**, MS^1^) and within the standard error range for such large ions.^44^ Once histone proteoforms were ejected, they were isolated and further fragmented. Additional HCD fragmentation causes protein ions to fragment along the backbone in predictable ion types (i.e., *b* and *y* ions, containing the N- and C-termini, respectively). These fragment ions were mapped onto the protein primary sequence for PTM localization on specific histones.^11^

**Supplemental Figure 4.**
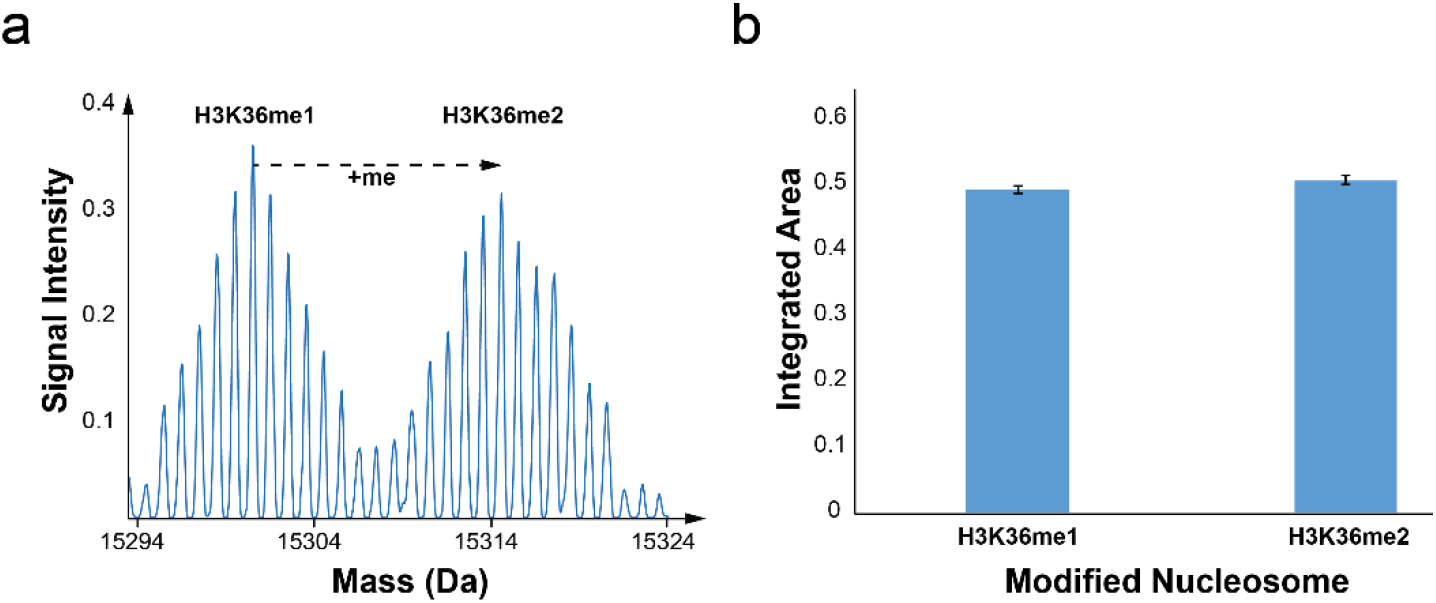
Nuc-MS analysis of equimolar H3K36me1 and H3K36me2 synthetic nucleosomes reveals equivalent signal intensities in the deconvoluted mass spectrum (**a**) and equivalent integrated areas upon quantitation (**b**), for three measurement replicates (n = 3). Specifically, the ratio of the integrated areas of ejected H3K36me1 and H3K36me2 proteoforms was calculated to be 49.2 ±2.5% and 50.8 ±3.3%, respectively.

**Supplemental Figure 5.**
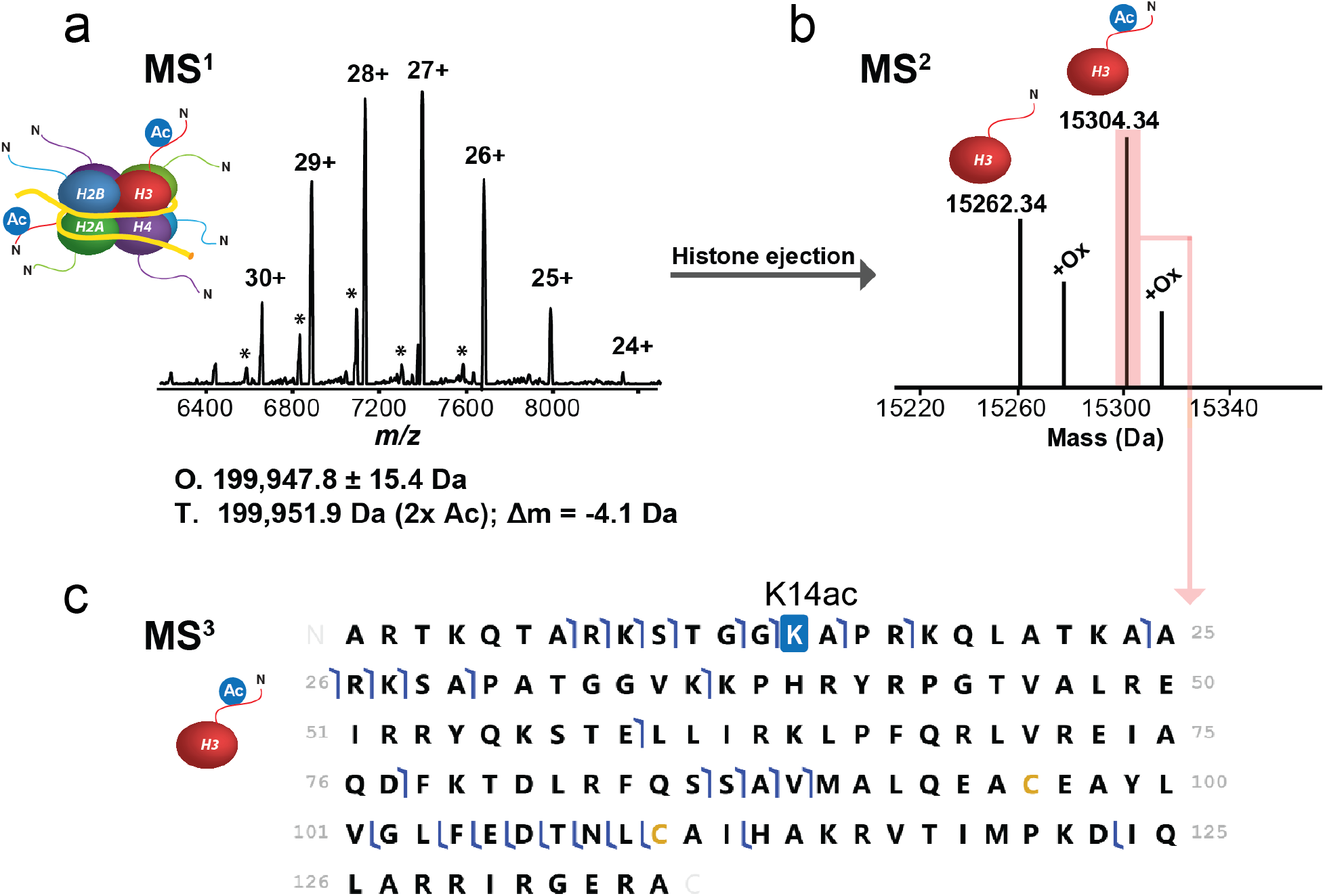
Nuc-MS of synthetic nucleosomes after treatment with PCAF acetyltransferase. **(a)** MS^1^: the full charge state distribution of a synthetic nucleosome acetylated with PCAF (492-658) *in vitro* (O, observed average mass and precision at 1σ; T, theoretical mass for doubly-acetylated nucleosome; Δ*m*, error). * Peaks denote a nucleosome species with a shorter DNA strand. **(b)** MS^2^: spectral region reporting monoisotopic mass values for histone H3 proteoforms ejected from acetylated nucleosomes in panel a. **(c)** MS^3^: graphical fragment map of acetylated H3, with complete localization of K14ac, consistent with the activity of PCAF (see supplemental discussion on this figure).^45^ Fragments were asserted and manually verified using TDValidator.

*Supplemental Discussion of in vitro nucleosome modification with PCAF.* The MS^1^ data on PCAF-treated nucleosomes yielded an observed average mass of 199,947.8 ± 15.4 Da, which is −4.1 Da from the theoretical mass of a doubly-acetylated nucleosome (**Supplemental Fig. 5a**). This mass error likely indicates an unresolved mixture of acetylated nucleosomes. MS^2^ thus helps to elucidate the nature of these acetylation events (**Supplemental Fig. 5b**). These data show masses corresponding to a mixture of 45% unmodified and 55% mono-acetylated H3, and also reveal significant oxidation. Thus, if we consider a mass addition from histone oxidation, the MS^1^ and MS^2^ data together suggest that the mixture consists of approximately equal amounts of doubly- and singly-acetylated nucleosomes, and up to 20% of unmodified nucleosomes.

*Finally*, the MS^3^ data can be used to localize the site of acetylations within H3. Hence, these data were converted into a fragmentation map revealing acetylation at K14, a proteoform consistent with PCAF activity^45^ (**Supplemental Fig. 5c**). Inspection of the data using TDValidator and mMass confirmed the K14ac site, and that no other positional acetylation isomer was present down to ∼5% relative abundance (when nucleosomes are treated for 5 minutes and for up to 2 hours). Previous studies of PCAF report acetylation at K9 and other sites beyond K14 when using H3 peptide substrates,^14^ but almost exclusive K14 acetylation when using nucleosomal substrate,^45, 46^ suggesting that substrate context yields differing results.

**Supplemental Figure 6.**
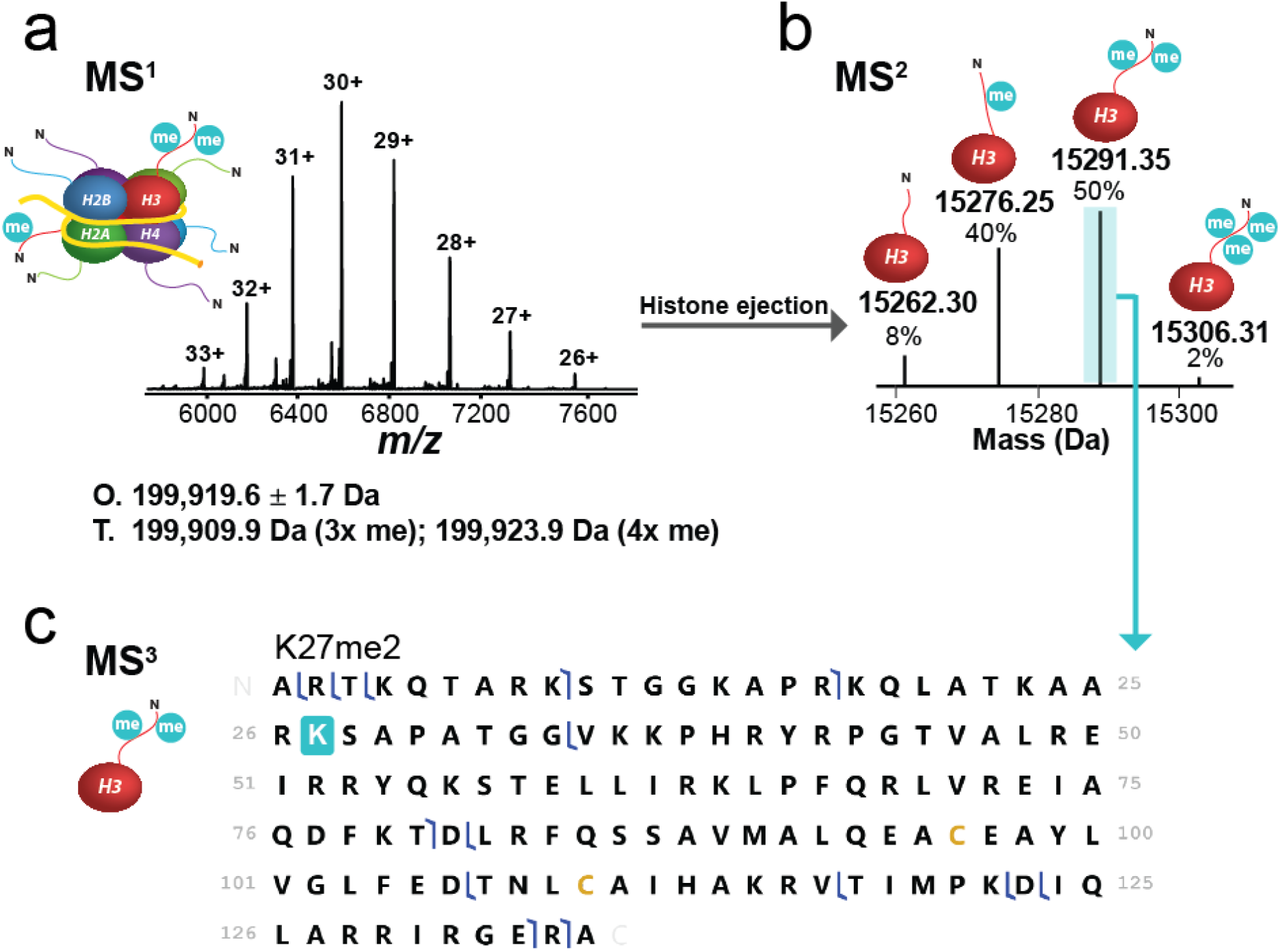
Nuc-MS data of synthetic nucleosomes upon treatment by PRC2 methyltransferase. **(a)** MS^1^: full charge state distribution of a synthetic nucleosome methylated with PRC2 *in vitro* (O, observed average mass and precision at 1σ; T, theoretical mass for tri- and tetra-methylated nucleosome). **(b)** MS^2^: spectral region reporting monoisotopic neutral masses for the methylated histone H3 proteoforms ejected from PRC2-trated nucleosomes. **(c)** MS^3^: graphical fragment map of di-methylated H3, consistent with H3.1K27me2.^47^ Fragments were asserted and manually verified using TDValidator.

*Supplemental Discussion of in vitro nucleosome modification with PRC2*. We analyzed synthetic nucleosomes modified by the Polycomb repressive complex 2 (PRC2), comprising full-length EZH2, SUZ12, EED, and RbAp46/48. The MS^1^ spectrum (**Supplemental Fig. 6a**) reports an observed average mass of 199,919.6 ±1.7 Da after PRC2 treatment, indicating a mixture of mostly tri- and tetra-methylated nucleosomes (199,909.9 Da and 199,923.9 Da, respectively). The ejected proteoforms in the MS^2^ (**Supplemental Fig. 6b**) reveal 8% unmodified, 40% mono-, 50% di-, and 2% tri-methylated H3. The low abundance of tri-methylation supports previous results that PRC2 is most efficient at catalyzing mono- and di-methylation.^15, 48^ Moreover, the MS^2^ data is consistent with a nucleosome mixture bearing on average three to four methylations. The fragmentation map created from the MS^3^ data of the di-methylated H3.1 proteoform (**Supplemental Fig. 6c**), is consistent with H3K27me2 installed by the SET domain of EZH2 within the PRC2 complex.

**Supplemental Figure 7.**
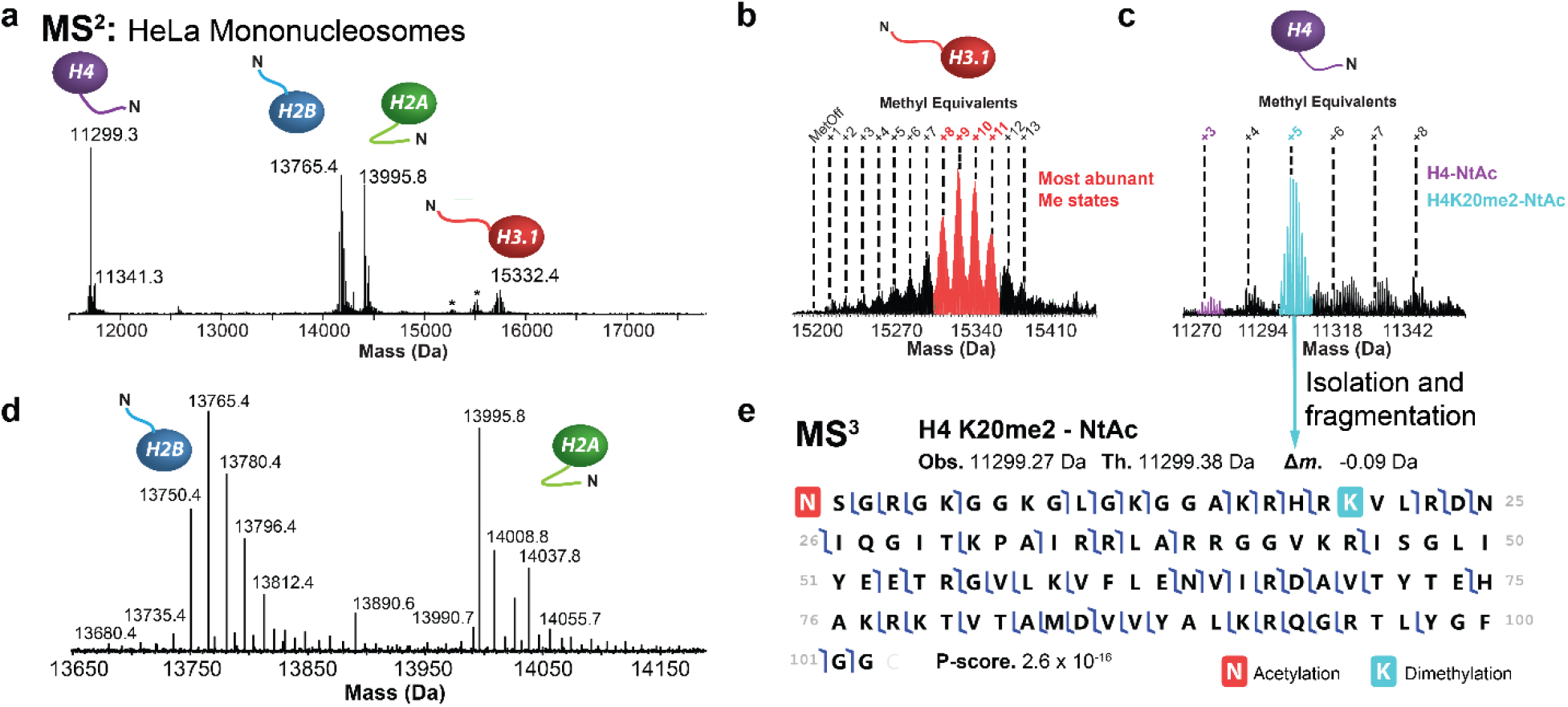
Nuc-MS analysis of endogenous mononucleosomes from HeLa. **(a)** MS^2^ spectrum of ejected histones (reported as deconvoluted, monoisotopic neutral masses), demonstrating detection of all core histones and their proteoform distributions; asterisks denote proteoform distributions consistent with N-terminal loss of the first two (AR) and four (ARTK) amino acids from H3.1. **(b-d)** Spectral regions in the mass domain containing the isotopic distributions for the proteoforms of histones H3.1, H4, and the monoisotopic masses of H2B and H2A proteoforms. **(e)** The proteoform highlighted in cyan in (c) was isolated and fragmented. The fragmentation data were then used to produce the fragmentation map shown, thus characterizing the proteoform as N-terminally acetylated H4K20me2.

**Supplemental Figure 8.**
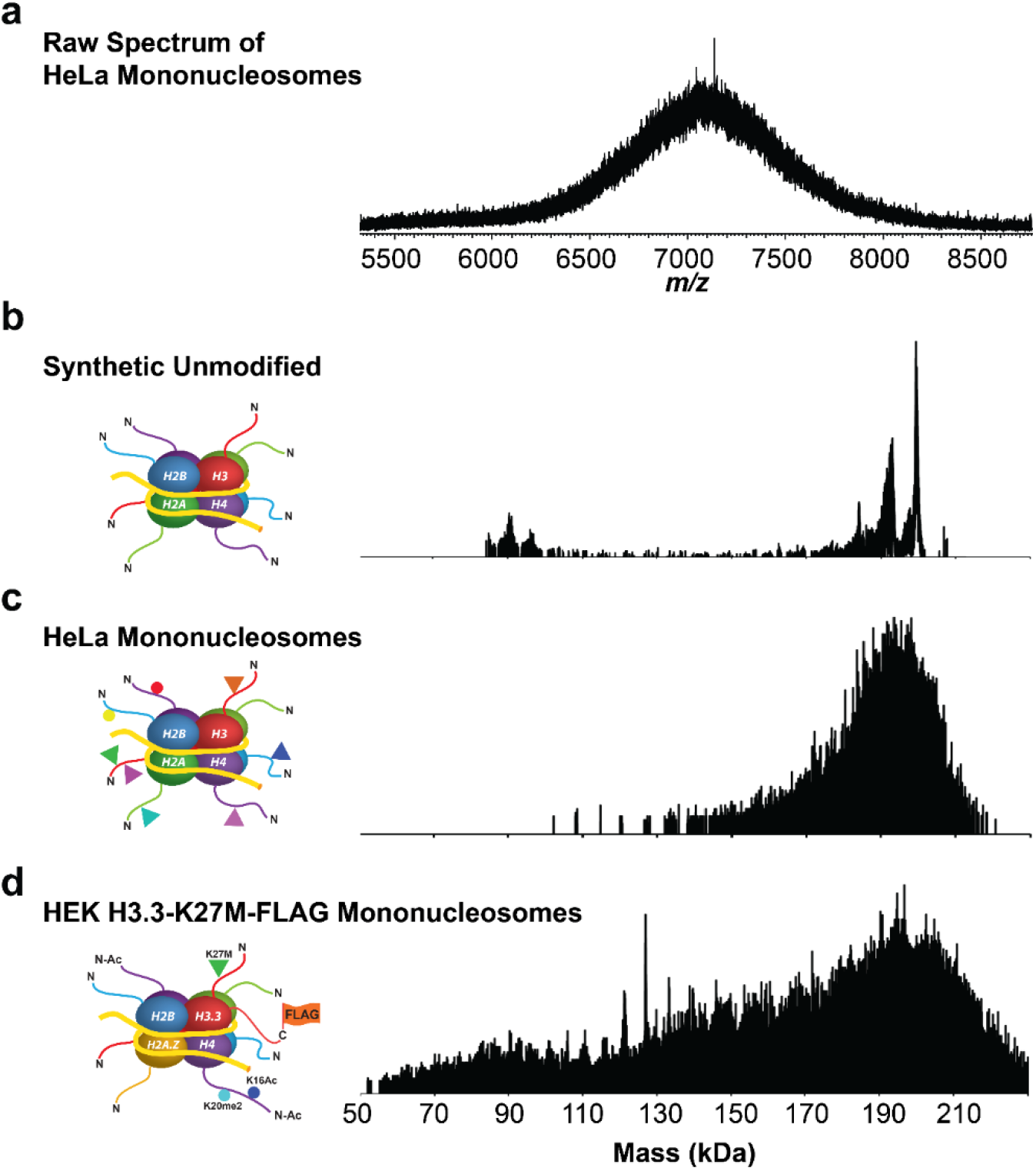
**(a)** Intact native mass spectrum of endogenous mononucleosomes from HeLa after MNase treatment.^21^ Given the chemical complexity of modifications present on the population of ∼100M nucleosomes contained in diploid cells, the native mass spectrum did not contain visible charge states needed for deconvolution and mass assignment. This motivated the use of the new approach of individual ion MS (I^2^MS), as described in the main text and methods. This novel analytical method relies on the direct assignment of charge to each individual mononucleosome ion (mainly found to be 26-28+ for the ∼1 million individual ions sampled to generate these spectra).^21^ **(b)** Mass-domain spectrum of synthetic, unmodified nucleosomes showing the predominant mass centered at 200 kDa. The smaller species correspond to nucleosomes with smaller lengths of DNA that likely result from PCR error during DNA synthesis. The largest, low abundance peak present is an artifact that stems from charge state mis-assignment. **(c)** Mass-domain spectrum of HeLa mononucleosomes, showing a mass distribution centered between 190-200 kDa. The apparent heterogeneity – a broad mass range between 170-210 kDa – likely stems from variability in MNase digestion products, resulting in mononucleosomes with varying lengths of DNA. **(d)** Mass-domain spectrum of H3.3K27M-FLAG mononucleosomes showing a broad distribution of masses, with the most abundant species centered around 200 kDa.

**Supplemental Figure 9.**
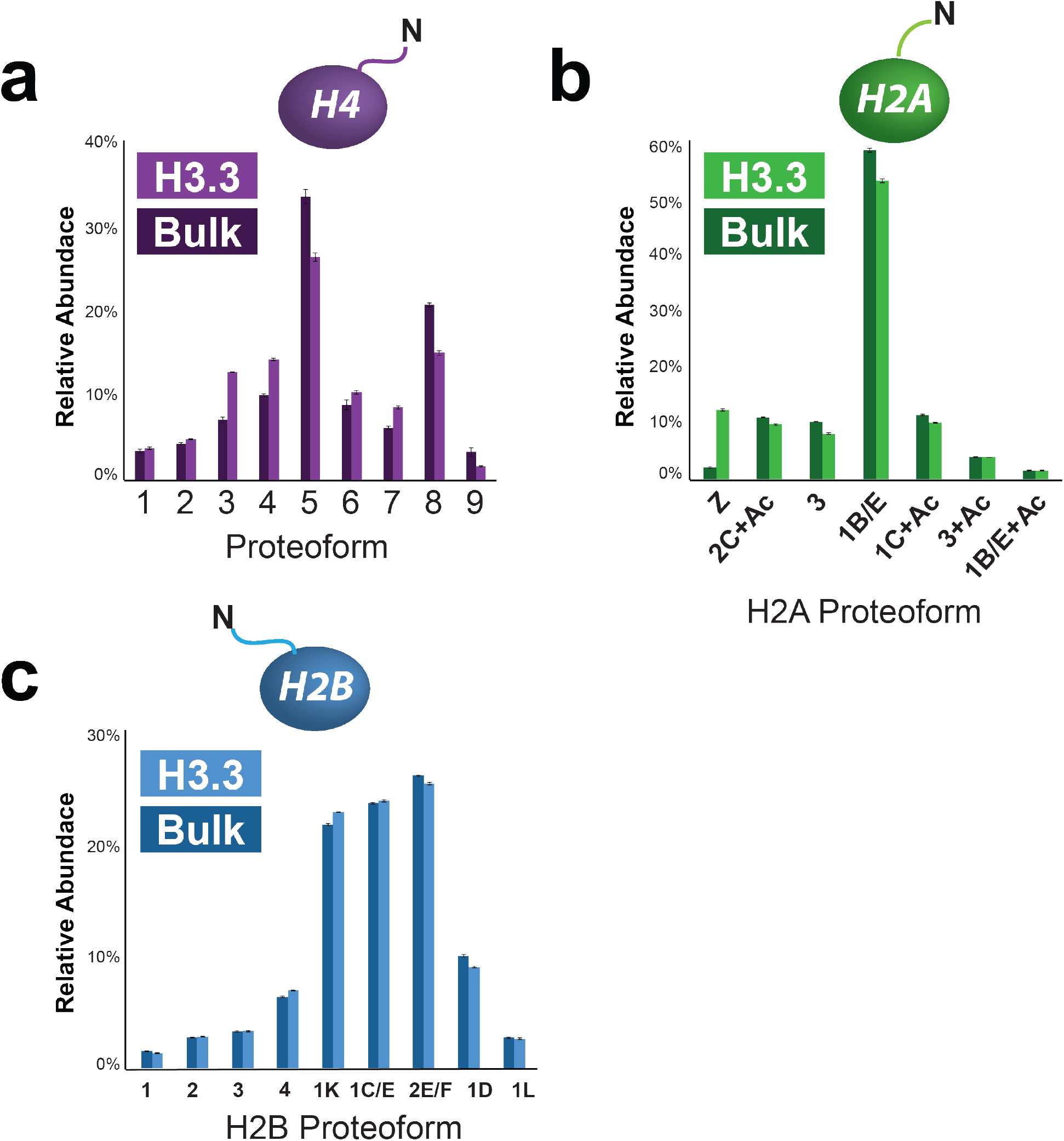
Quantitation of **(a)** H4, **(b)** H2A, and (**c**) H2B proteoforms ejected from HEK bulk nucleosomes (Bulk) compared to those enriched for H3.3-FLAG (H3.3). The sum of the abundances of all proteoforms in each histogram equals 100%. H4 proteoforms are displayed in order of methyl equivalence and are listed from low to high mass as proteoforms 1-9, as shown in **Fig. 3c**. The following are important H4 proteoforms mentioned in the main text: H4-NtAc (3), H4K20me1-NtAc (4), H4K20me2-NtAc (5), and H4K20me1Ac-NtAc (7).

**Supplemental Figure 10.**
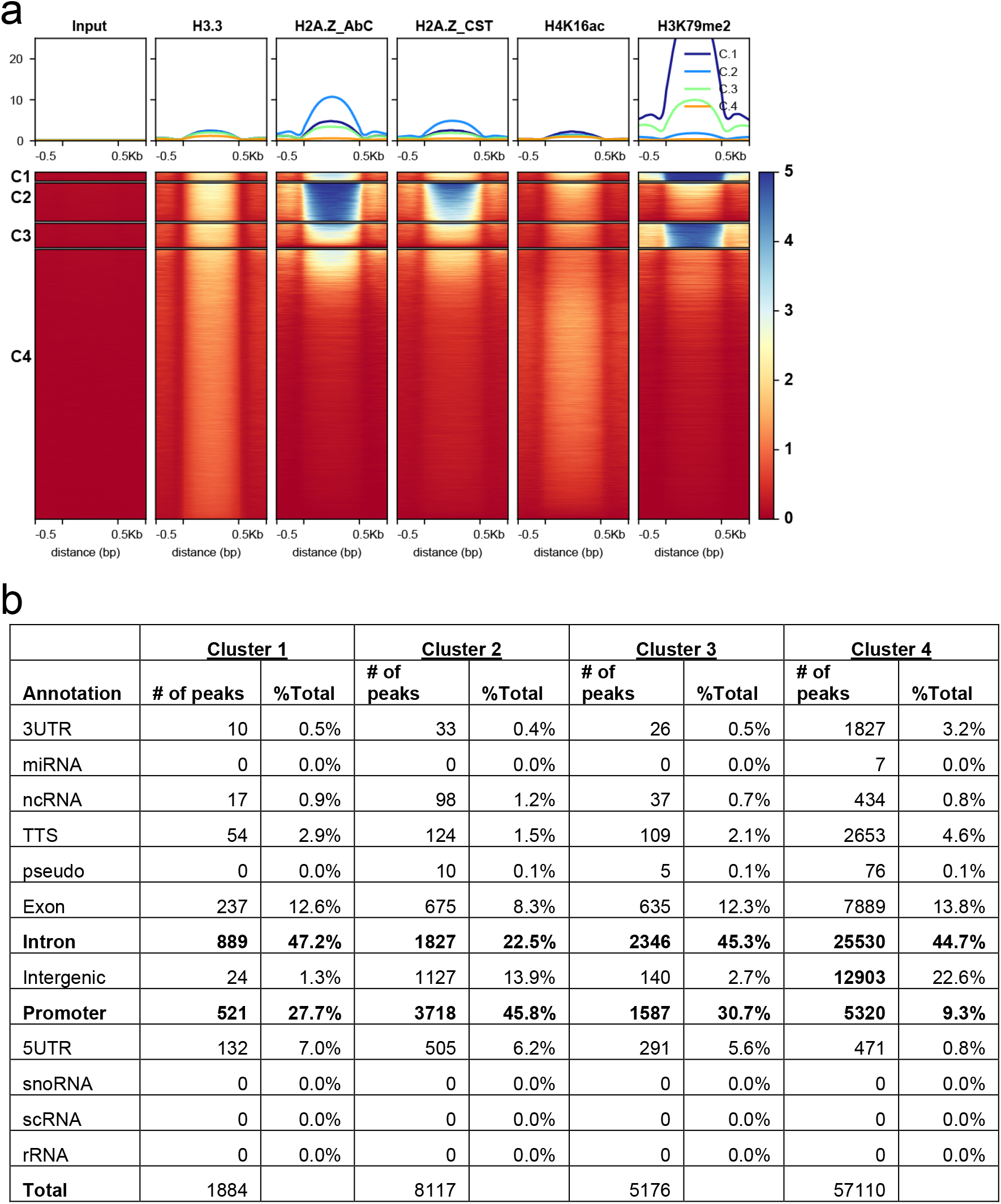
**(a)** Heatmap centered on H3.3 peaks ± 0.5 kb showing the correlation of ChIP-seq signal for input, H3.3, H3K79me2, H2A.Z (using antibodies from Abcam and Cell Signaling Technologies, CST), and H4K16ac, and extending 0.5 kb in each direction. **(b)** Table summarizing genetic composition of each cluster listed in **(a)**.

**Supplemental Figure 11.**
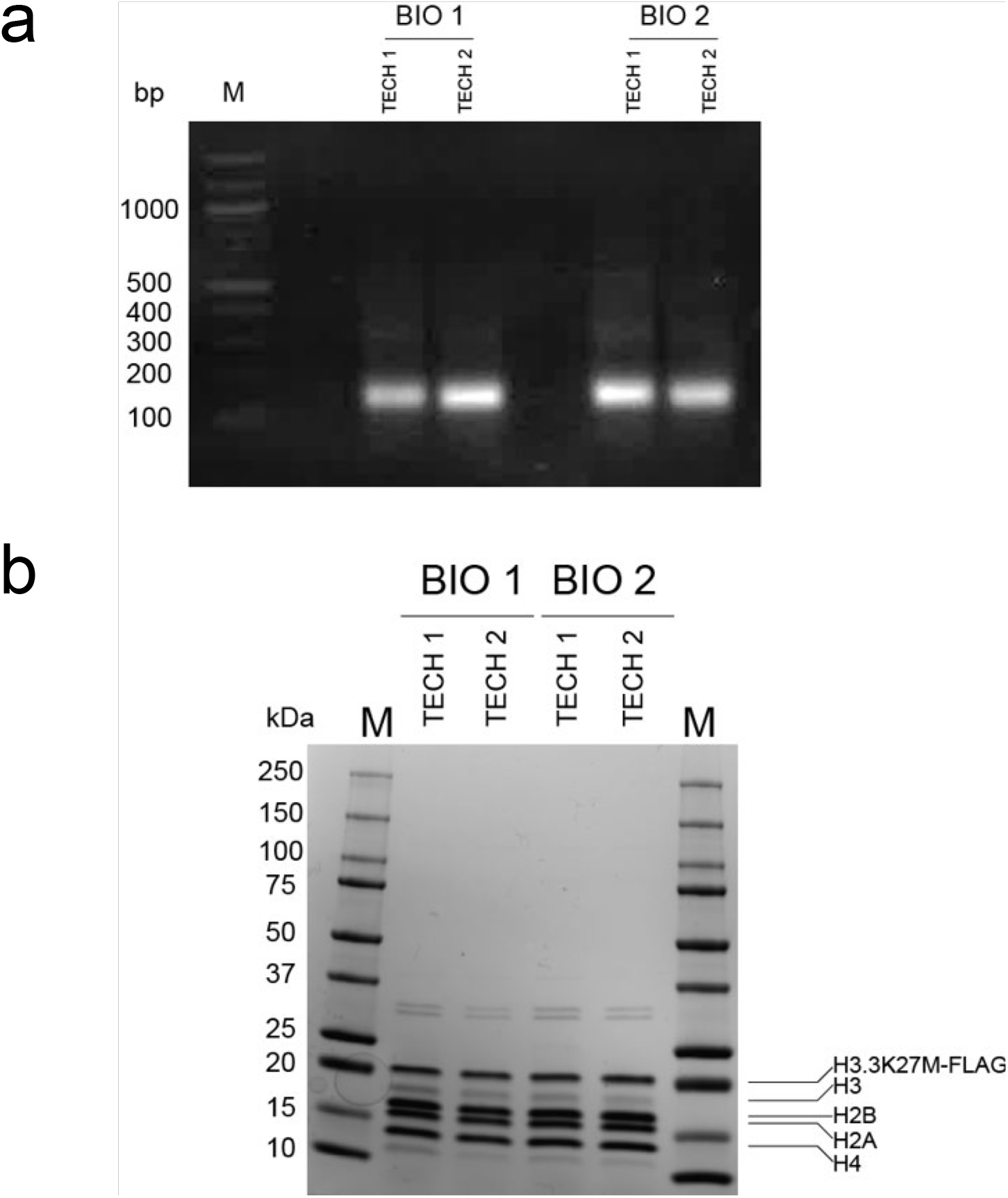
DNA and protein gels showing masses consistent with FLAG-tagged H3.3K27M mononucleosomes **(a)** and its constituent histone proteoforms **(b)**.

**Supplemental Figure 12.**
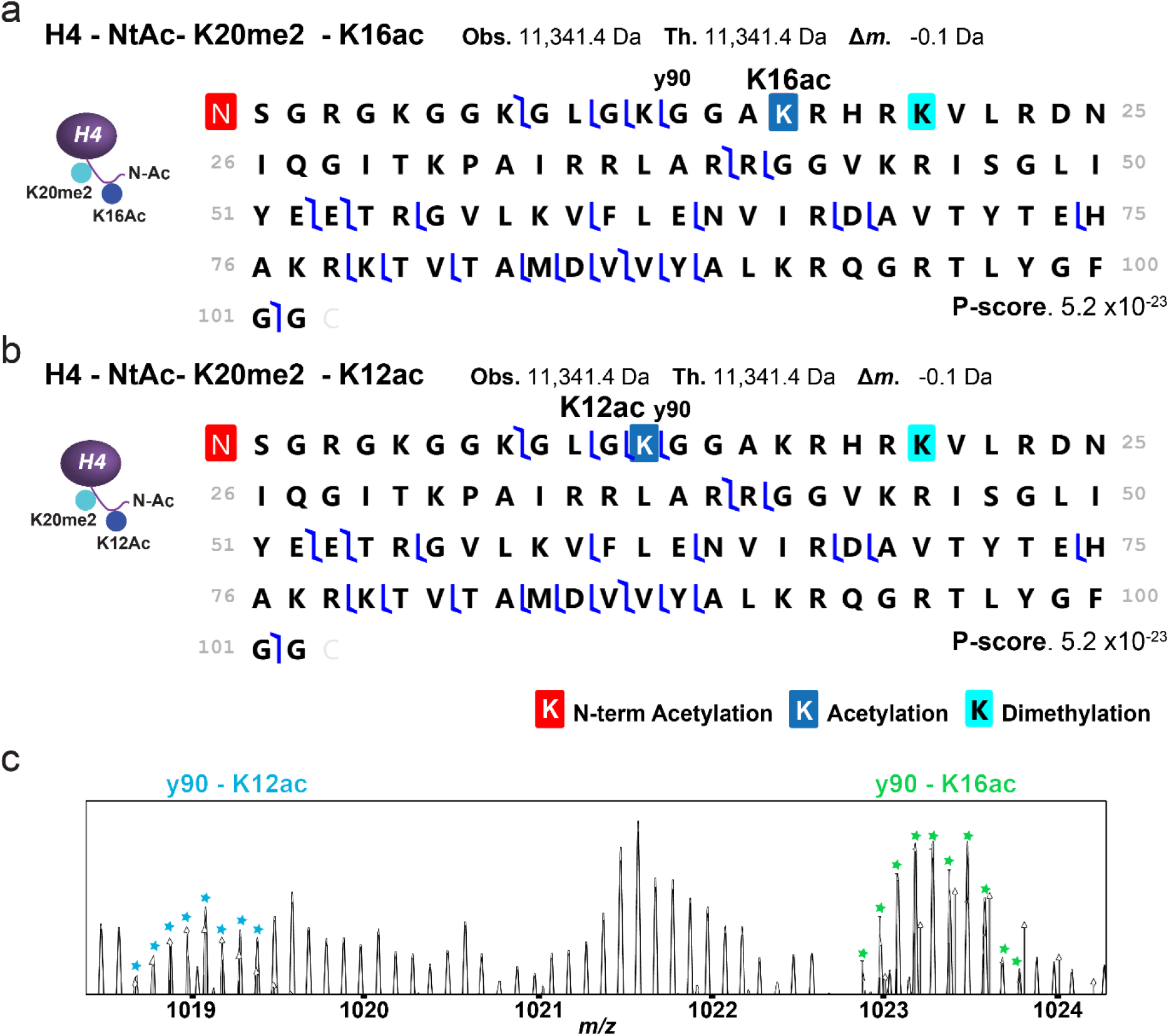
Fragmentation maps of H4 proteoforms detected from H3.3K27M mononucleosomes. Fragmentation data support H4K20me2 co-existing with K16 and K12 acetylation (**a** and **b**, respectively). Moreover, diagnostic ion y90 reveals that H4K16ac is the predominant proteoform, as shown by the isotopic distributions in the TDValidator output^42^ of panel **(c)**.

**Supplemental Figure 13.**
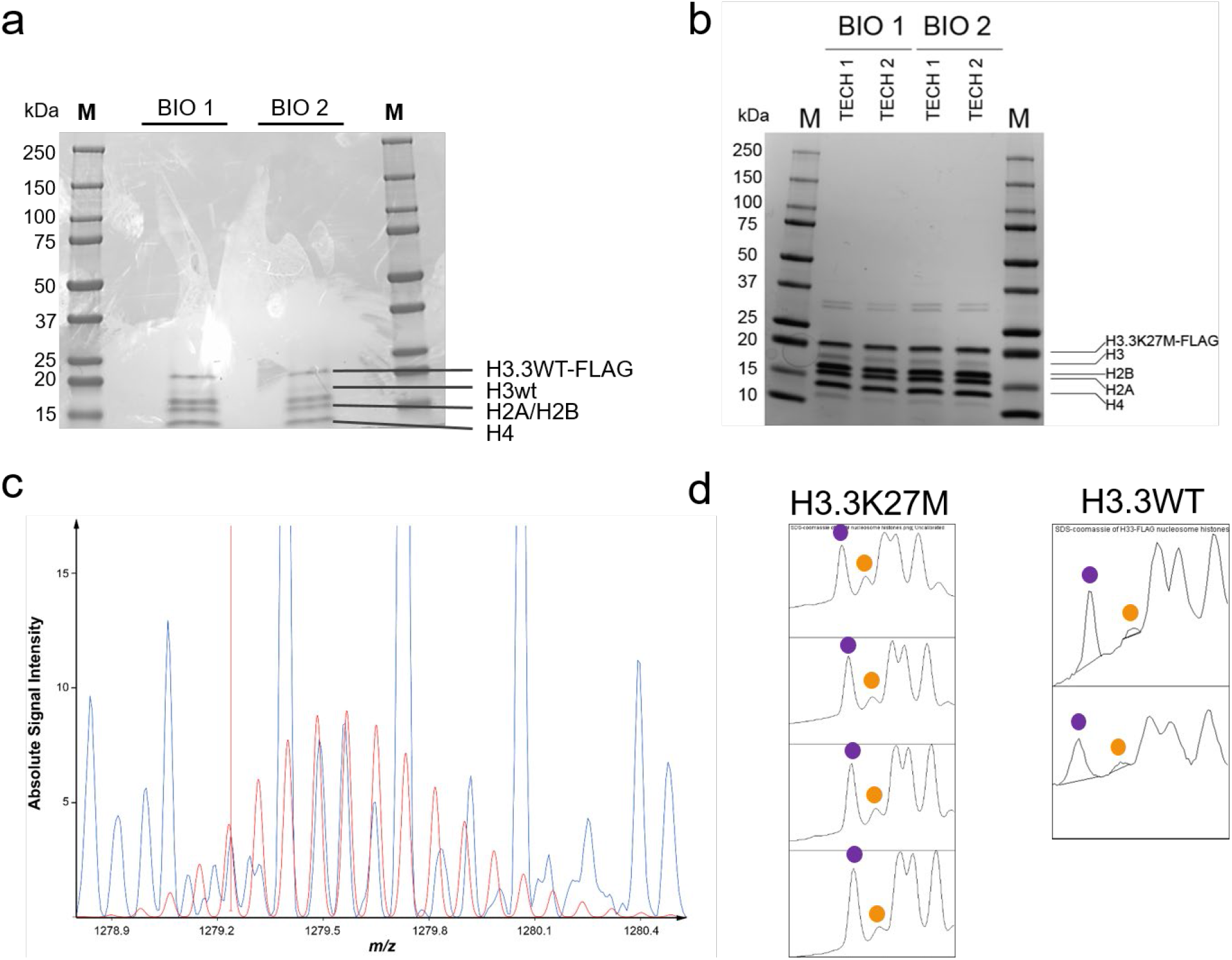
Coomassie-stained SDS-PAGE of histones enriched after FLAG IP of (**a**) H3.3WT (**b**) H3.3K27M nucleosomes, showing significantly lower abundance of H3 WT relative to the FLAG-tagged counterpart. (**c**) Theoretical isotopic distribution (red) overlaid on the experimental isotopic distribution for the 12+ charge state of H3.1WT with 5 methyl equivalents derived from K27M nucleosomes, reflecting the low intensity of this proteoform distribution. (**d**) ImageJ-generated traces of the intensities of the Coomassie-stained bands in the gels of **(a)** and **(b)**. The areas underneath the purple-labeled peaks (H3.3K27M-FLAG or H3.3WT-FLAG) and orange-labeled peaks (H3WT) were integrated to determine the relative ratios of these proteoforms. These were found to be 88% H3.3K27M-FLAG compared to 12% H3WT (n = 4), and 90% H3.3WT-FLAG compared to 10% H3WT (n = 2).

**Supplemental Figure 14.**
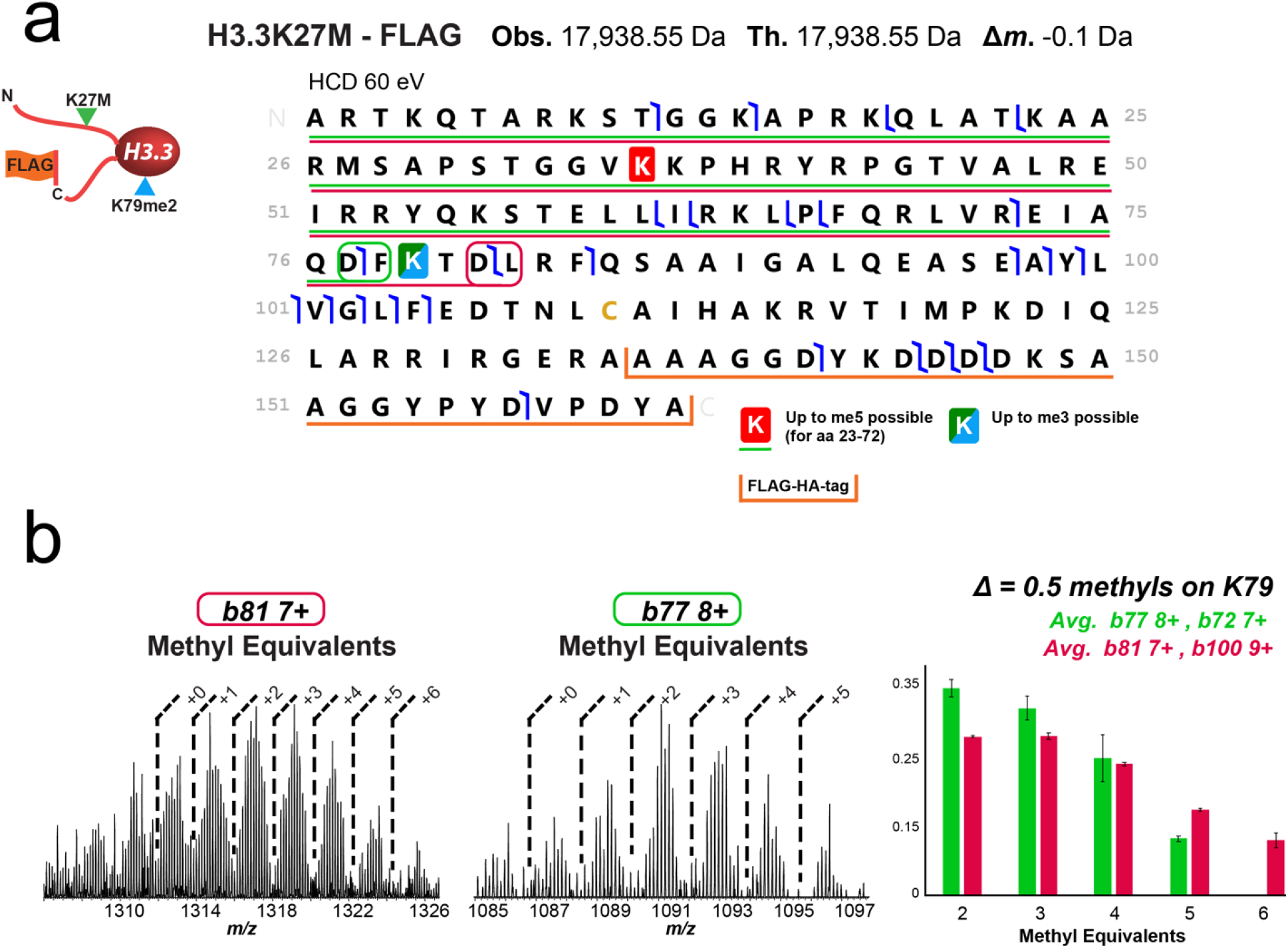
Fragmentation of H3.3K27M by MS^3^ at HCD of 60 eV. **(a)** Analysis of backbone fragment ions of H3.3K27M proteoform enables partial localization of methylation. Even though fragmentation data are not sufficient to precisely localize methylations between residues 23-72, the data are consistent with two to three methyl equivalents in this region. **(b)** Subtracting the average methyl equivalence for diagnostic ions *b81* and *b100* (red) from that of *b77* and *b72* (green), results in an average decrease of 0.5 methyl equivalents when moving from the *b81* to *b77* fragment ion pair. This indicates a 50% probability of mono-methylation and 25% probability of di-methylation between amino acids 78-81 (FKTD); K79 being the most likely residue to contain this PTM. A 25% probability of K79me2 constitutes at least ∼15-fold enrichment relative to bulk H3 (1-2%). Additional fragment ions (*y100-101* 8+, *y96* 8+ & 9+, and *y97* 9+) further localize the assignment of K79me2 being co-occurring with H3.3K27M and part of the reprogrammed code on these mutant nucleosomes. The data do not support PTMs N-terminal to H3K23, indicating that these are low abundance and below the limit of detection.

**Supplemental Figure 15.**
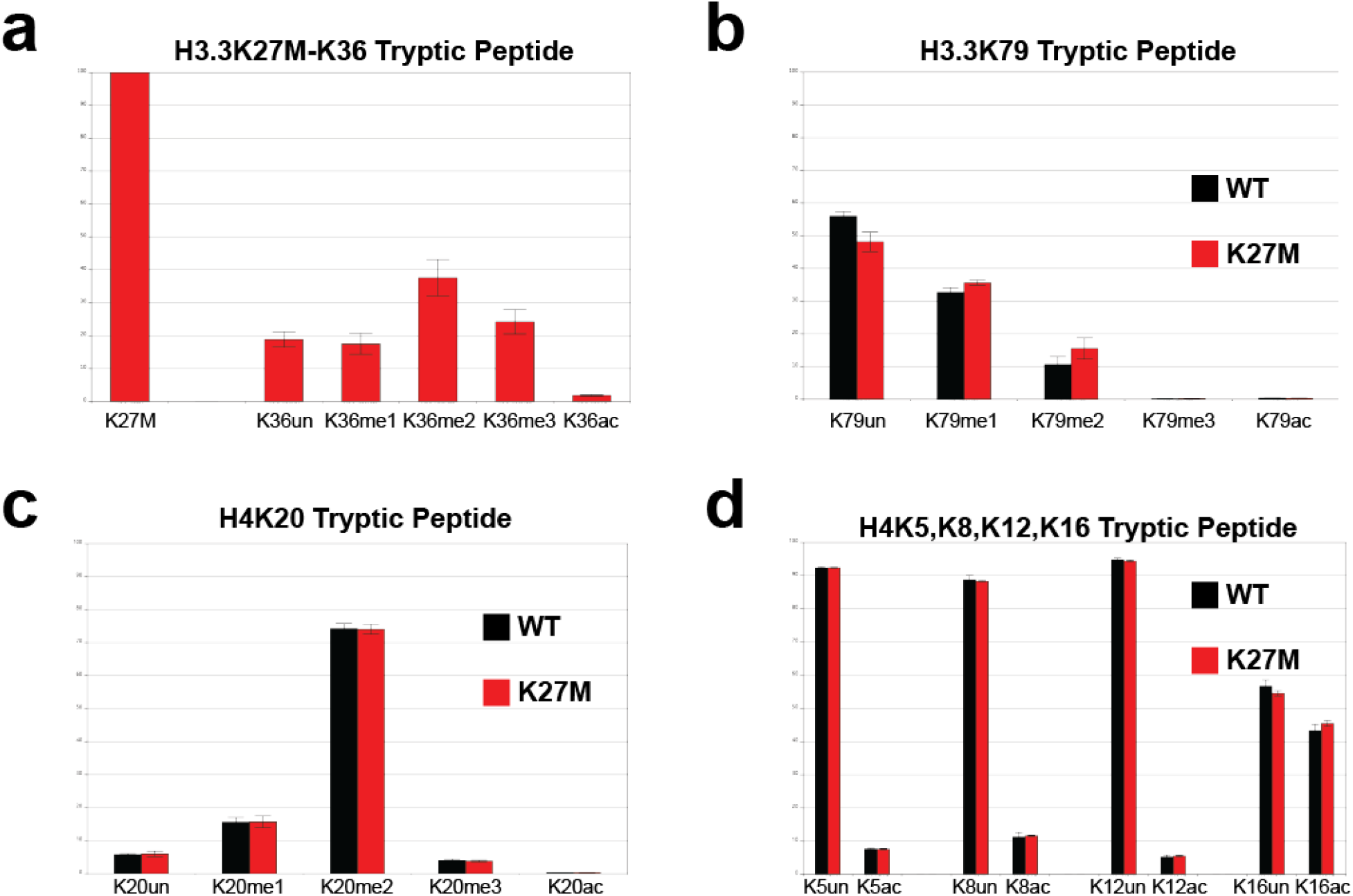
Analysis of tryptic peptides of histones from bulk chromatin of HCT116 cells supports detection at the proteoform levels made by the new Nuc-MS workflow, including: (**a)** H3.3K36me2, **(b)** H3.3K79me2, **(c)** H4K20me2, and **(d)** acetylated H4 proteoforms.

**Supplemental Figure 16.**
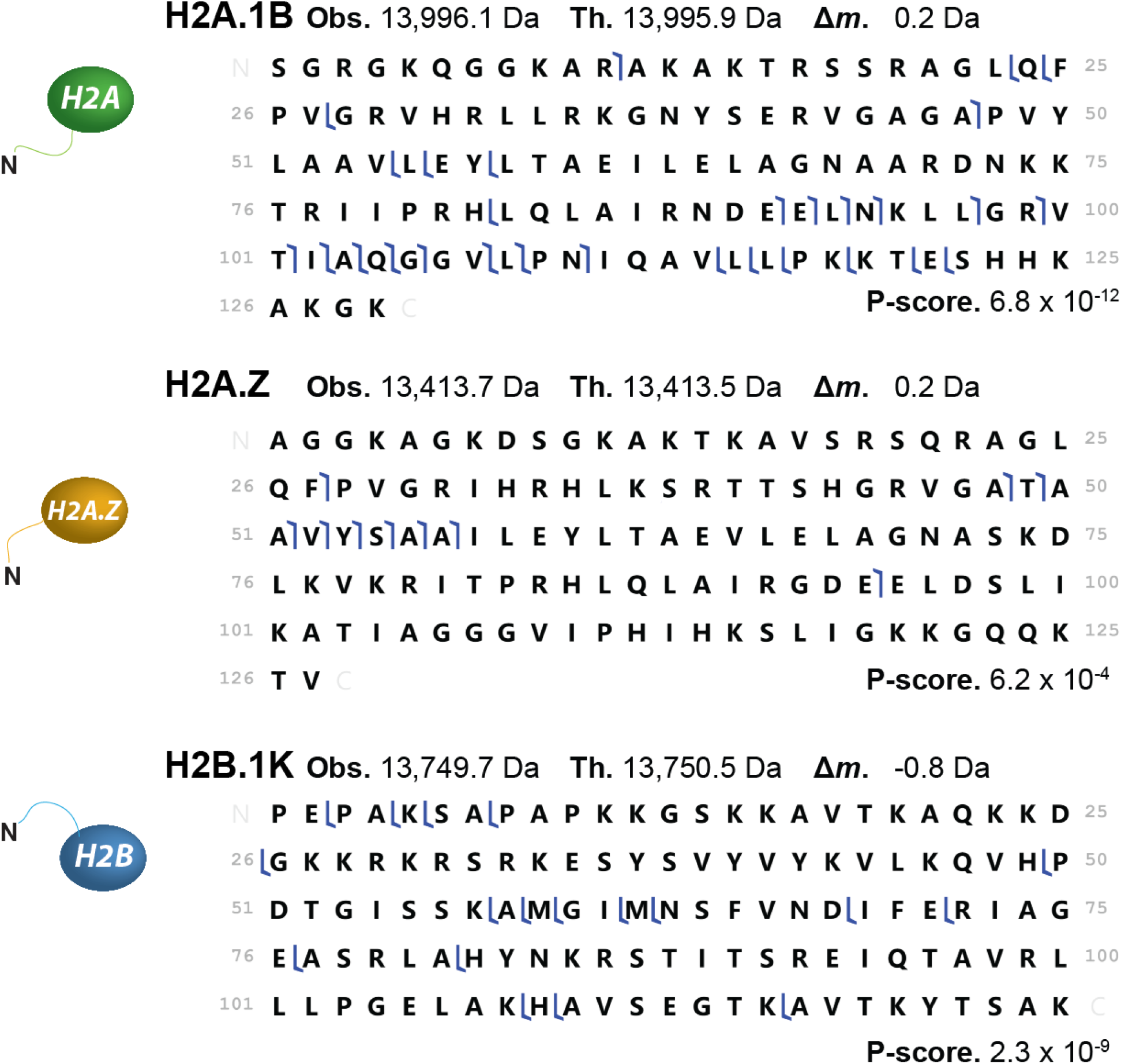
Fragmentation maps showing identification of H2A, H2A.Z, and H2B proteoforms after ejection from intact H3.3K27M mononucleosomes by Nuc-MS. Fragments were manually validated using TDValidator. P-scores were calculated using ProSight Lite.

**Supplemental Figure 17.**
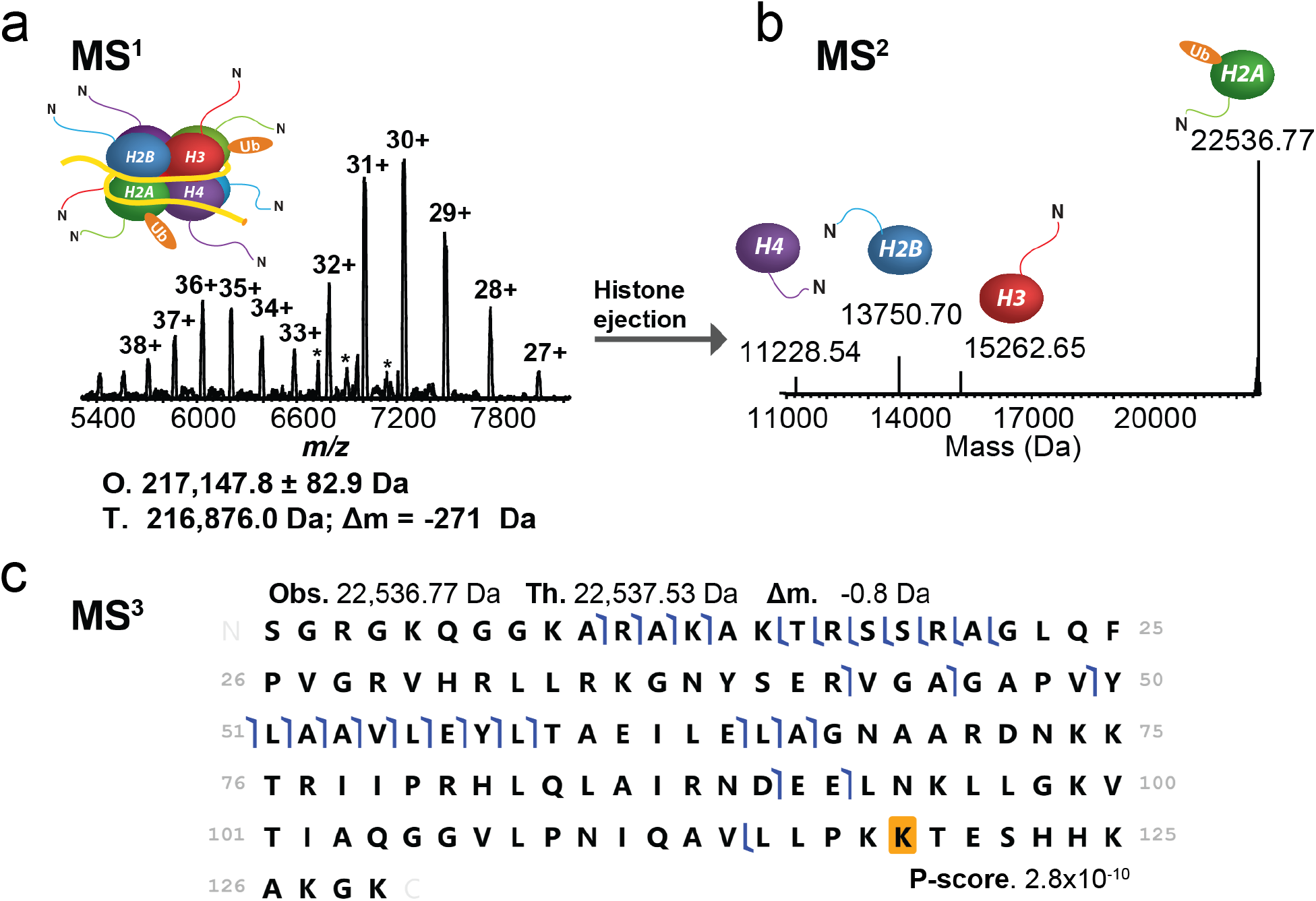
Nuc-MS data of a synthetic nucleosome ubiquitinated at H2AK119 serves as a positive control to detect H2A ubiquitination in endogenous samples. **(a)** MS^1^: full charge state distribution of a synthetic mono-ubiquitinated nucleosome (O, observed average mass and precision at 1σ; T, theoretical mass for a homotypic nucleosome carrying two mono-ubiquitins on H2AK119). **(b)** MS^2^: spectral region reporting monoisotopic neutral masses for the ubiquitinated histone H2A and ejected proteoforms of the other core histones. **(c)** MS^3^: graphical fragment map of ubiquitinated H2A, consistent with mono-ubiquitination at lysine 119. Fragments were manually validated using TDValidator. P-scores were calculated using ProSight Lite.

*Supplemental Discussion on ubiquitinated nucleosomes.* Given the potential for cross-talk between H2A, H2B and H3.3 in euchromatic regions, we hypothesized that there might be changes in ubiquitination in K27M nucleosomes.^33^ As a positive control, we analyzed mono-ubiquitinated nucleosomes created *in vitro* and found that the new method efficiently detects this large PTM at both the nucleosome and histone levels; however, we did not detect ubiquitinated histones in the endogenous samples. Therefore, we have confidence in the sensitivity of Nuc-MS and can assert that H2A or H2B ubiquitination does not occur on >1% of H3.3WT or K27M nucleosomes in any of the HEK293T cells we analyzed by Nuc-MS.

**Supplemental Figure 18.**
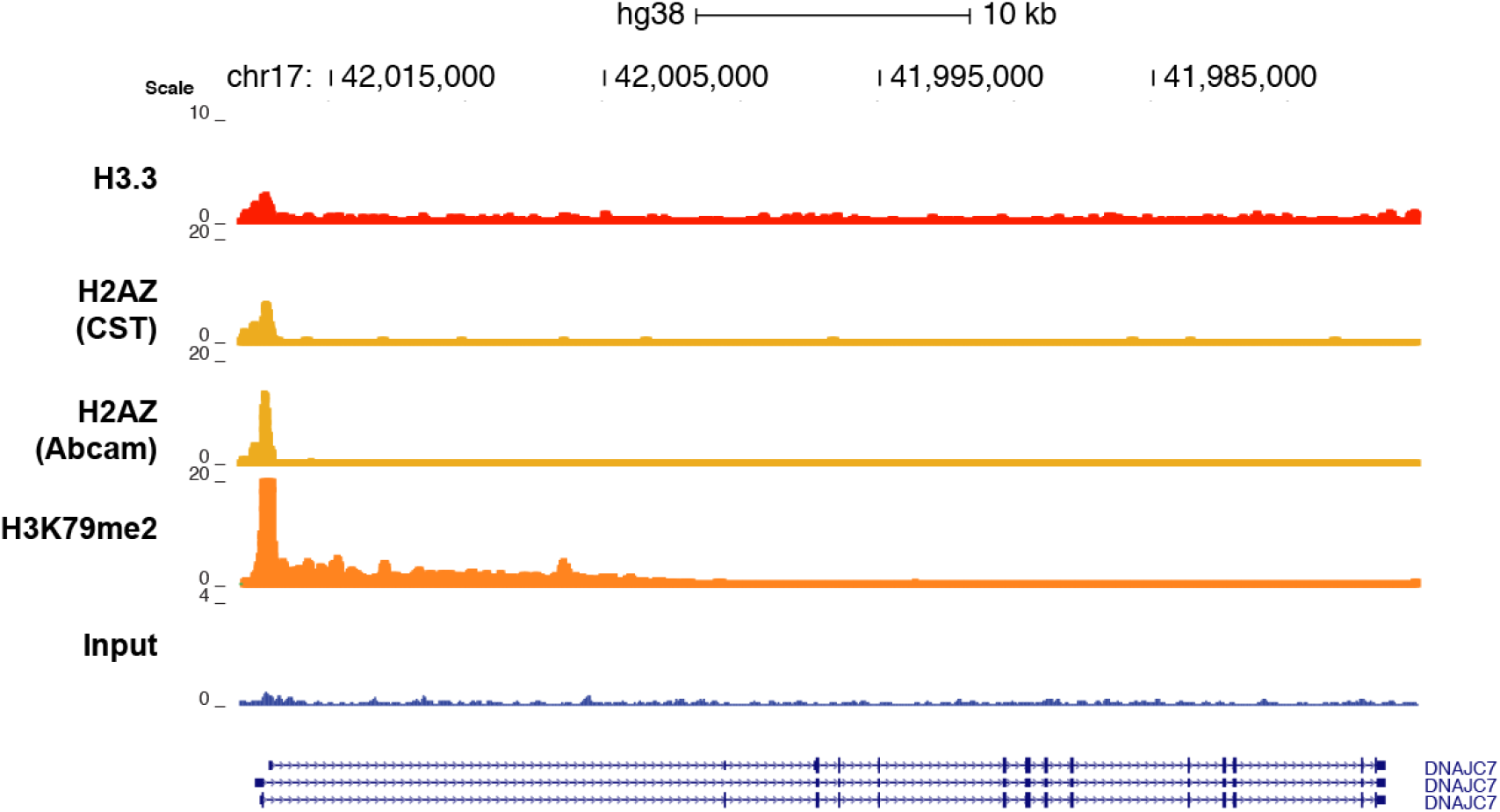
Zoom-in of a representative gene showing peak overlap for H3.3, H2A.Z and H3K79me2. Moreover, the H3K79me2 track shows that the K79me2 mark is propagated into the gene body, unlike H3.3 and H2A.Z signals, which are confined to the transcription start site.

**Supplemental Figure 19.**
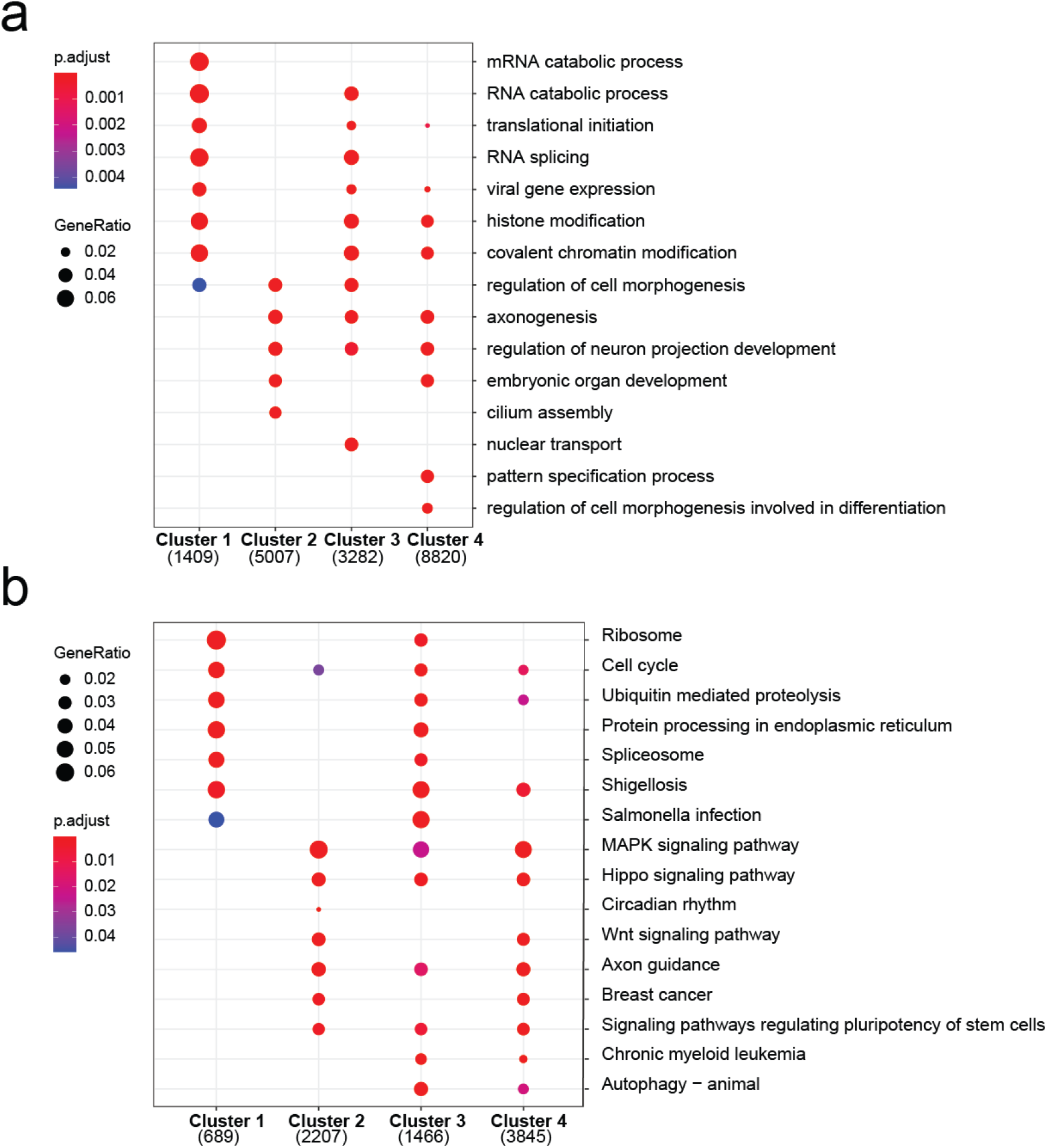
Gene composition of each cluster presented in the ChIP-seq heatmaps in **Figs. 2** and **3** and **Supplemental Fig. 10**, described by the gene ontology terms **(a)** biological process or **(b)** molecular function.

**Supplemental Table 1.** Calculated *p*-values for abundance differences between histone proteoforms derived from bulk chromatin of HEK and H3.3-FLAG enriched nucleosomes. Red *p*-values denote statistically significant differences for a given proteoform’s intensity between the samples with Bonferroni-corrected α-values.

| Histone | Proteoform | Bonferroni-corrected α-value | $p$-value |
| --- | --- | --- | --- |
| **H4** | 1 | 0.00556 | 0.0926 |
|  | 2 | 0.00556 | **0.00352** |
|  | 3 | 0.00556 | **7.49E-06** |
|  | 4 | 0.00556 | **2.34E-06** |
|  | 5 | 0.00556 | **2.15E-04** |
|  | 6 | 0.00556 | 0.0139 |
|  | 7 | 0.00556 | **6.81E-05** |
|  | 8 | 0.00556 | **1.00E-05** |
|  | 9 | 0.00556 | **0.00360** |
| **H2A** | Z | 0.00714 | **2.71E-07** |

**Supplemental Table 2.** Calculated *p*-values for abundance differences between histone proteoforms derived from H3.3WT and H3.3K27M nucleosomes. Red *p*-values denote statistically significant differences for a given proteoform’s intensity between WT and K27M samples with Bonferroni-corrected α-values.

| Histone | Proteoform | Bonferroni-corrected α-value | *p*-value |
| --- | --- | --- | --- |
| **H4** | 1 | 0.00556 | 0.0327 |
|  | 2 | 0.00556 | 0.0316 |
|  | 3 | 0.00556 | **0.00309** |
|  | 4 | 0.00556 | 0.01582 |
|  | 5 | 0.00556 | **0.00119** |
|  | 6 | 0.00556 | 0.0264 |
|  | 7 | 0.00556 | 0.0163 |
|  | 8 | 0.00556 | **1.01E-04** |
|  | 9 | 0.00556 | **0.00106** |
| **H3.3** | 1 methyl equivalent | 0.00714 | **0.00435** |
|  | 2 methyl equivalents | 0.00714 | **5.76E-04** |
|  | 3 methyl equivalents | 0.00714 | **0.00426** |
|  | 4 methyl equivalents | 0.00714 | 0.0648 |
|  | 5 methyl equivalents | 0.00714 | **0.00284** |
|  | 6 methyl equivalents | 0.00714 | **1.93E-05** |
|  | 7 methyl equivalents | 0.00714 | 0.836 |
| **H2A** | Z | 0.00714 | **1.73E-4** |
|  | 2C+Ac | 0.00714 | 0.418 |
|  | 3 | 0.00714 | 0.526 |
|  | 1B/E | 0.00714 | 0.0661 |
|  | 1C+Ac | 0.00714 | **0.00143** |
|  | 3+Ac | 0.00714 | 0.655 |
|  | 1B/E+Ac | 0.00714 | 0.0952 |

## Supplemental References for ChIP-seq methods

-Andrews S. (2010). FastQC: a quality control tool for high throughput sequence data. Available online at: http://www.bioinformatics.babraham.ac.uk/projects/fastqc

-Heinz S, Benner C, Spann N, Bertolino E et al. Simple Combinations of Lineage-Determining Transcription Factors Prime cis-Regulatory Elements Required for Macrophage and B Cell Identities. Mol Cell 2010 May 28;38(4):576-589.

-Langmead, B., Trapnell, C., Pop, M., and Salzberg, S.L. (2009). Ultrafast and memory-efficient alignment of short DNA sequences to the human genome. Genome Biol 10, R25.

-Lee et al., 2006. Nature Protocols. “Chromatin immunoprecipitation and microarray-based analysis of protein location”

-Ramírez, Fidel, Devon P. Ryan, Björn Grüning, Vivek Bhardwaj, Fabian Kilpert, Andreas S. Richter, Steffen Heyne, Friederike Dündar, and Thomas Manke. deepTools2: A next Generation Web Server for Deep-Sequencing Data Analysis. Nucleic Acids Research (2016).

-Ross-Innes CS, Stark R, Teschendorff AE, Holmes KA, Ali HR, Dunning MJ, Brown GD, Gojis O, Ellis IO, Green AR, Ali S, Chin S, Palmieri C, Caldas C, Carroll JS (2012). “Differential oestrogen receptor binding is associated with clinical outcome in breast cancer.” Nature, 481, −4.

-Vo et al., 2017. Cell Reports. “Inactivation of Ezh2 Upregulates Gfi1 and Drives Aggressive Myc-Driven Group 3 Medulloblastoma”

-Yu G, Wang L, Han Y, He Q (2012). “clusterProfiler: an R package for comparing biological themes among gene clusters.” OMICS: A Journal of Integrative Biology, 16(5), 284-287.

-Zhang, Y., Liu, T., Meyer, C.A., Eeckhoute, J., Johnson, D.S., Bernstein, B.E., Nussbaum, C., Myers, R.M., Brown, M., Li, W., et al. (2008). Model-based Analysis of ChIP-Seq (MACS). Genome Biol 9, R137.

